# Lateral hypothalamic input engages a disinhibitory microcircuit in the dorsal raphe to promote behavior activation

**DOI:** 10.64898/2026.02.26.708342

**Authors:** Renata Sadretdinova, Arturo Marroquin Rivera, Zakaria Benmammar, Maryse Pinel, Vincent Émond, Chenqi Zhao, Frédéric Calon, Benoit Labonté, Christophe D. Proulx

## Abstract

Behavioral activation involves coordination between hypothalamic and brainstem systems that regulate movement and internal state, but the circuit logic underlying this interaction remains poorly defined. The dorsal raphe nucleus (DRN), a major serotonergic hub, integrates diverse inputs to influence behavioral inhibition and activation, yet how hypothalamic signals shape its activity is unclear. Here, we identify a disinhibitory pathway through which the lateral hypothalamus (LHA) promotes behavioral activation via DRN. Using intersectional viral tracing, electrophysiology, and single-nucleus RNA sequencing, we show that LHA inputs preferentially target GABAergic DRN neurons that locally inhibit 5-HT neurons. Silencing these DRN neurons innervated by LHA increased locomotor and repetitive behaviors, decreased local inhibition, and enhanced cFos activity in serotonergic neurons, consistent with circuit-level disinhibition. Molecular profiling revealed distinct transmitter identities and transcriptional signatures of LHA-targeted versus LHA-projecting DRN populations. Together, these findings delineate a hypothalamic–raphe circuit that transforms hypothalamic drive into serotonergic activation, revealing a mechanism by which the lateral hypothalamus promotes behavioral activation through local inhibitory control.

**HIGHLIGHTS:**

- LHA inputs preferentially target non-serotonergic, transcriptionally distinct DRN neurons
- LHA-innervated DRN neurons form extensive local inhibition
- Silencing LHA-innervated DRN neurons reduced local inhibitory tone and activate serotonin neurons
- Disrupting this circuit drives behavioral activation with repetitive motor pattern, not anxiety

## INTRODUCTION

Serotonin (5-HT) neurons are highly heterogeneous in both function and connectivity^1,2^. The dorsal raphe nucleus (DRN), the principal source of forebrain serotonin, has long been implicated in diverse sensorimotor, cognitive, and affective processes^3,4^. Classically, serotonergic activity has been associated with behavioral inhibition and impulse control: reduced serotonergic signaling increases impulsivity^5^, whereas activation of DRN 5-HT neurons promotes patience in delayed-reward paradigms^6,7^. In contrast, other studies have demonstrated that enhancing serotonergic transmission, either pharmacologically or through direct activation of DRN 5-HT neurons, can increase mobility and active coping in aversive contexts such as the forced- or tail-suspension tests^8,9^. This apparent contradiction suggests that the behavioral consequences of 5-HT activity depend on context. For instance, population activity of DRN 5-HT neurons shows negative correlations with movement onset in neutral environments but positive correlations under stress^10^. When monitoring single-cell calcium activity, this variability is further reflected in the heterogeneity of DRN 5-HT neurons, with 38% showing positive correlation with locomotion and 18% showing negative correlation in the open field test^11^. Together, these findings suggest that 5-HT neurons do not uniformly inhibit or promote movement but instead exert context-dependent control shaped by their upstream inputs and local network interactions^1,2^.

Besides 5-HT neurons, the DRN also comprises GABAergic, glutamatergic, and dopaminergic cells^2^. Distinct long-range afferents from the forebrain, hypothalamic, and brainstem converge onto DRN 5-HT and GABAergic neurons, forming input-specific circuits. Notably, excitatory and inhibitory inputs from the same upstream structure can differentially target 5-HT versus GABAergic neurons, providing opposing influences on downstream targets^1^. These features imply that local inhibitory control within the DRN is a key node through which afferent signals shape serotonergic output and, consequently, behavioral state^1,12,13^. To determine how a defined upstream pathway engages this inhibitory microcircuit, it is therefore essential to dissect DRN regulation at the level of input- and output-defined populations^1,14^.

The lateral hypothalamus (LHA) is one of the major long-range inputs to the DRN^14–16^. The LHA is a functionally heterogeneous hub that integrates internal state signals to coordinate behavioral outputs^17–20^. Activation of GABAergic LHA neurons increases locomotor activity and can elicit compulsive-like behaviors, whereas their inhibition or ablation suppresses spontaneous movement^21–23^. More specifically, GABAergic LHA projections to the ventral tegmental area (VTA) promote reinforcement and behavior activation by disinhibiting dopamine neurons through local inhibition of VTA GABAergic interneurons^24^. In contrast, glutamatergic LHA neurons projecting to the VTA drive aversive responses, defensive behaviors, and avoidance^24–26^. Given its established role in regulating goal-directed behaviors, the LHA is well positioned to modulate DRN circuits that engage the context-appropriate locomotor activity.

Although the DRN is strongly interconnected with the LHA, the anatomical and functional organization of this pathway remains poorly characterized. The LHA sends excitatory and inhibitory projections to serotonergic and non-serotonergic neurons in the DRN^12,27^. Yet, the cellular composition and synaptic logic through which these inputs engage distinct DRN subpopulations to influence behavior are still unclear. Recent work from our group showed that activity of LHA axon terminals in DRN coincides with 5-HT neurons activation and movement onset in both the open field and tail-suspension test, and that optogenetic stimulation of LHA→DRN terminals promotes locomotion and active coping^27^. However, how excitatory and inhibitory inputs from the LHA are integrated within the DRN microcircuits to regulate serotonergic activity and behavior remains unknown.

In this study, we combine viral-tracing, single-nucleus RNA sequencing, electrophysiology, and behavioral assays to dissect the organization and function of the LHA→DRN pathway. We show that LHA axons preferentially target non-serotonergic GABAergic neurons in the DRN, providing both excitatory and inhibitory inputs that converge to produce an indirect activation of serotonergic neurons through local inhibition. Chronic silencing of DRN neurons innervated by the LHA increases locomotion and active coping behaviors without affecting anxiety or social interactions. Together, these findings reveal an input-specific disinhibitory microcircuit through which hypothalamic signals engage serotonergic output to promote behavioral activation.

## RESULTS

### Reciprocal LHA-DRN connectivity is organized through largely distinct, non-serotonergic populations

The DRN contains molecularly and functionally heterogeneous neuronal populations with complex input-output organization^2^. To identify DRN neurons receiving monosynaptic input from the LHA, we injected an AAV1-Cre viral vector, capable of anterograde transsynaptic transport^28^, and a Cre-dependent AAV-DIO-eGFP into the DRN, enabling selective labeling of DRN neurons postsynaptic of LHA axons (hereafter referred to as DRN^:LHA^ neurons; **Figure 1A**). In parallel, to label DRN neurons projecting back to LHA (DRN→LHA), we co-injected a retrograde AAV expressing FlpO (retroAAV-flpO)^29^ into the LHA together with an FlpO-dependent AAV-fDIO-mCherry into the DRN. This intersectional strategy allowed direct comparison of input- and output-defined DRN populations while accounting for possible retrograde labeling by AAV1.

**Figure 1.**
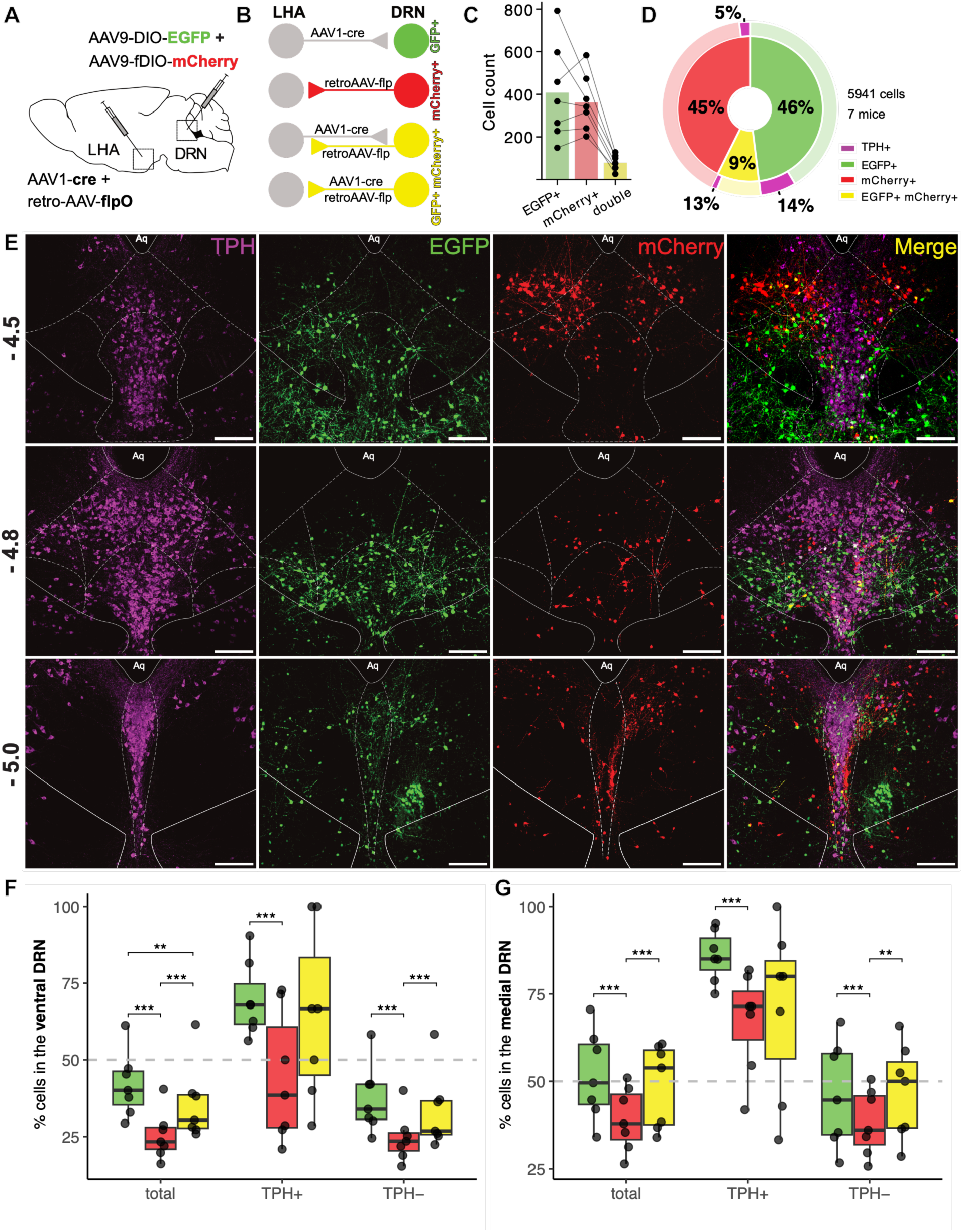
Reciprocal LHA-DRN connectivity is organized through largely distinct, non-serotonergic populations. A. Schematic of the viral strategy. eGFP was expressed in the cells receiving synaptic input from the DRN; mCherry was expressed in DRN cells projecting onto the LHA. B. Schematic illustration of four theoretical labeling outcomes of our viral strategy. C. Total number of eGFP+, mCherry+, and double-labeled eGFP+ mCherry+ DRN neurons across mice (n = 7 mice; 5,941 cells quantified). D. Proportions of eGFP+, mCherry+, and double-labeled eGFP+ mCherry+ neurons within the DRN (n = 7 mice). The fraction of TPH+ neurons within each categories is indicated in magenta. E. Representative coronal sections of the DRN showing TPH immunostaining (magenta), GFP (green), and mCherry (red). Sections are shown at rostral (−4.5), medial (−4.8), and caudal (−5.0) levels relative to bregma. Scale bar: 200 µm. F. Percentage of eGFP+, mCherry+, and eGFP+ mCherry+ neurons in the ventral DRN, relative to the total number of neurons in both dorsal and ventral DRN. G. Percentage of eGFP+, mCherry+, and eGFP+ mCherry+ neurons in the medial DRN, relative to the total number of neurons in both medial and lateral DRN. Boxplots show the median and interquartile range (IQR), with whiskers extending to 1.5× IQR; points represent individual mice. Statistical comparison in (F) and (G) were performed using binomial generalized linear mixed models (GLMM) with random intercept for each mouse followed by Tukey-adjusted pairwise comparisons. \*\**p* < 0.01, and \*\*\**p* < 0.001.

This approach labeled DRN neurons receiving LHA input (eGFP+), DRN neurons projecting to the LHA (mCherry+), and a smaller subset of neurons expressing both fluorophores (**Figure 1B**). Double-labeled neurons may reflect either dual viral retrograde labeling or a limited population participating in reciprocal LHA-DRN connectivity (**Figure 1B**).

Across seven mice, we identified a total of 5,941 labeled DRN neurons. Of these, 46% ± 8.0 were eGFP+ (408 ± 226 cells per animal), 44,4% ± 7.0 were mCherry+ (361 ± 132 cells per animal), and 9,4% ± 3.0 were double-labeled (79 ± 36 cells per animal; **Figure 1C-D**). Double-labeled neurons represented 17% ± 6.0 of eGFP+ cells and 18% ± 4.0 of mCherry+ cells, indicating limited overlap between DRN neurons receiving LHA input and those projecting back to LHA.

Immunostaining for tryptophane hydroxylase 2 (TPH) revealed that serotonergic neurons constituted a minority of all labeled populations. Specifically, 14% ± 4 of eGFP+, 5% ± 3 of mCherry+, and 13% ± 8 of doubled-labeled neurons were TPH+ (**Figure 1D**), indicating that both LHA-innervated and LHA-projecting DRN populations are predominantly non-serotonergic.

Because retrograde viral vectors can exhibit cell-type-dependent tropism^29^, we asked whether the low fraction of serotonergic neurons observed among LHA-connected DRN populations (∼5%) reflected true circuit organization or technical limitations of viral labeling. To address this, we validated our findings using alternative approaches. Injection of AAV11, which has been reported to display broader retrograde tropism^30^, similarly resulted in limited infection of TPH+ neurons (>5%) and yielded colocalization rate between GFP and mCherry (10% ± 4%) comparable to those obtained with retroAAV **(Figure S1A and S1B)**. In this condition, double-labeled neurons accounted for 22% ± 5% of eGFP+ neurons and 17% ± 8% of mCherry+ neurons.

In contrast, the classical retrograde tracer cholera toxin B (CTb) robustly labeled serotonergic neurons, with 21% ± 7% of CTb-positive neurons coexpressing TPH **(Figure S1C and S1D)**. Notably, despite this increased labeling of 5-HT neurons, colocalization between CTb and eGFP remained low (12% ± 3%), comparable to that observed using viral tracers. Together, these controls indicate that the limited overlap between LHA-innervated and LHA-projecting DRN neurons does not primarily reflect viral tropism, but instead supports the conclusion that reciprocal LHA-DRN connectivity involves largely distinct neuronal populations, with serotonergic neurons representing a minority of each group.

We next examined the spatial distribution of these populations along the dorsal-ventral and medial-lateral axes of the DRN using atlas-aligned anatomical landmarks. eGFP+ neurons were relatively evenly distributed, with 42% (95% CI: 35-49) located ventrally and 51% (95% CI: 43-58) medially **(Figure 1F and 1G)**. In contrast, mCherry+ neurons were biased towards the dorsolateral DRN, with only 25% (95% CI: 20-30) located ventrally and 39% (95% CI: 32-47) medially. Double-labeled neurons exhibited an intermediated distribution with 38% (95% CI: 20-43) located ventrally and 49% (95% CI: 41-58) medially.

When serotonergic and non-serotonergic populations were analyzed separately, TPH+ eGFP+ neurons were predominantly located in the ventromedial DRN. In contrast, TPH+ mCherry+ neurons remained largely medial but were more evenly distributed along the dorso-ventral axis.. Non-serotonergic eGFP+ neurons were enriched in dorsomedial regions, whereas non-serotonergic mCherry+ neurons were more dorsolaterally distributed.

Occasional labeling was observed in regions adjacent to the DRN, including the midbrain reticular tegmental nucleus (MRtN) and dorsal tegmental nucleus (DTN), which were excluded from all quantitative analyses. Because the MRtN contains GABAergic neurons that project to DRN 5-HT neurons^14^, injection volumes were reduced in subsequent functional experiments to minimize off-target labeling.

Together, these data indicate that LHA axons predominantly innervate non-serotonergic DRN neurons and that DRN receiving LHA input are largely distinct from those projecting to the LHA, revealing an anatomically segregated organization of reciprocal LHA-DRN connectivity.

### RNA sequencing reveals the transcriptional profile of LHA-targeted DRN neurons

To molecularly characterize DRN neurons defined by their anatomical connectivity with the LHA, we performed single-nucleus RNA sequencing (snRNA-seq) using a modified Vector-seq strategy that enabled detection of viral transgene transcript. DRN neurons were labeled using the same virus strategy described in **Figure 1**, allowing direct transcriptional profiling of DRN neurons receiving LHA inputs (eGFP+) and those projecting to the LHA (mCherry+) within the same samples (**Figure 2A)**.

**Figure 2.**
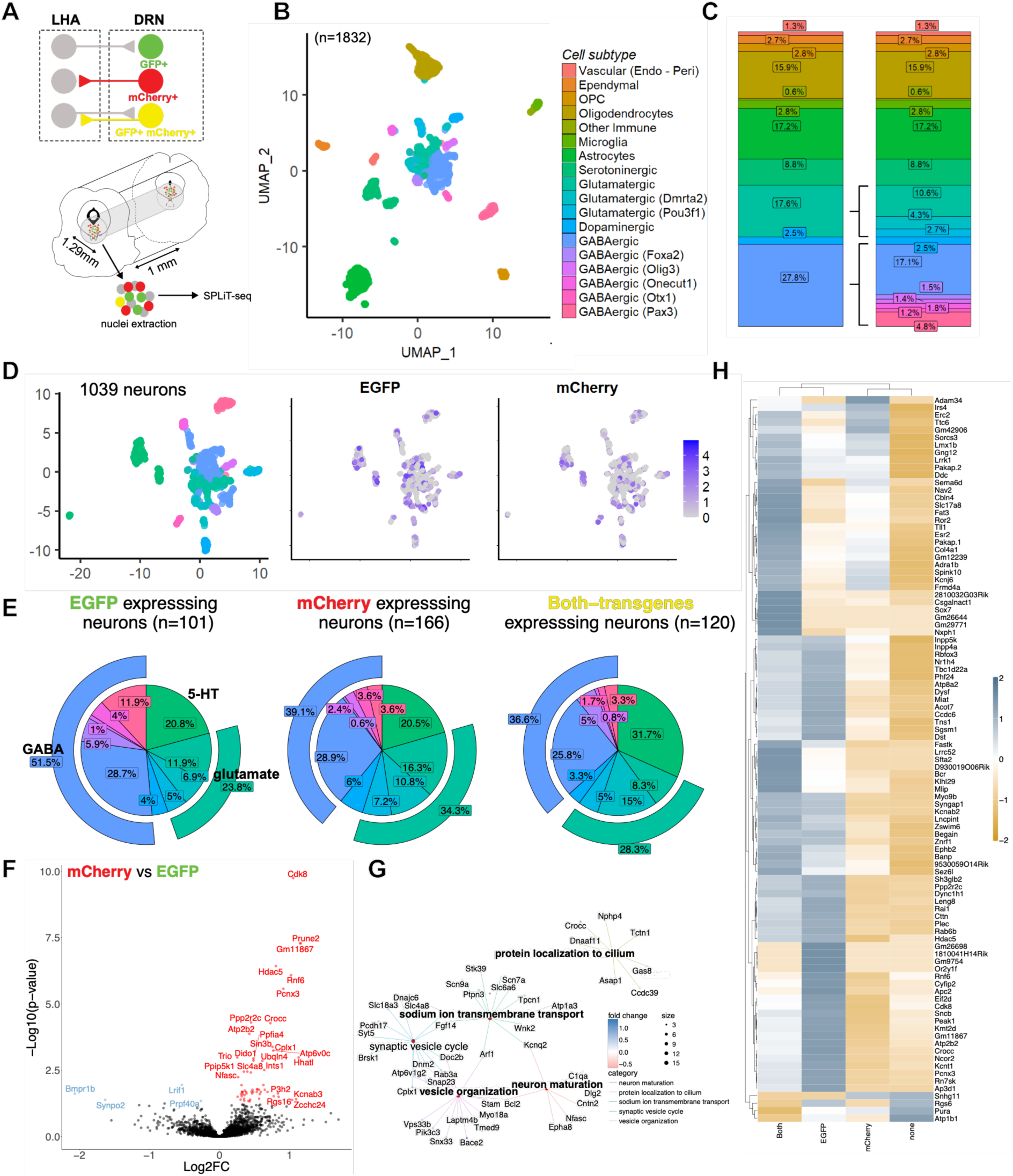
RNA sequencing reveals the transcriptional profile of LHA-targeted DRN neurons. A. Schematic of the experimental workflow. DRN tissue containing virally labeled neurons was microdissect, nuclei were isolated, and single-nucleus RNA sequencing was performed using SPLiT-seq. Viral transgene transcripts were detected to identified DRN neurons receiving LHA inputs (eGFP+), projecting to the LHA (mCherry+), or expressing both transgenes. B. UMAP projection of the 1832 nuclei included in the dataset. The cell types were annotated using MapMyCells followed by manual curation based on canonical marker expression. C. Proportion of major cell types and neuronal subtypes in the whole dataset. D. UMAP projection of the 1039 annotated neuronal nuclei. Expression level of eGFP and mCherry transcripts herein evaluated are overlaid on the same UMAP projection and the expression corresponds to the normalized counts. E. Pie charts of the distribution of neuronal subclasses among eGFP+ (n = 101), mCherry (n = 166), and double-labeled (n = 120). Transgene-positive cells were defined as ≥5 transgene reads per nucleus. F. Volcano plot showing the differentially expressed genes (DEGs) between eGFP+ and mCherry+ neurons identified using the GLM model and its corresponding Wald test for the β-coefficient (FDR-adjusted). Red points indicate genes enriched in mCherry+ neurons; blue points indicate genes enriched in eGFP+ neurons. G. Cnetplot of the top five enriched Gene Ontology (GO) terms associated with DEGs in F. H. Heatmap showing the average normalized expression of the top DEGs identified by Wilcoxon Rank-sum tests comparing each transgene-defined group (eGFP+, mCherry, double-labeled, none) against all other neuronal nuclei.

Unsupervised clustering and manual annotation based on canonical markers^31^ identified a cellular composition dominated by neurons (56.7%), mostly represented by GABAergic (27.8%) and glutamatergic (17.6%) neuronal subtypes (**Figure 2B-C and S2A-B**). Among the non-neuronal populations, astrocytes (17.2%) and oligodendrocytes (15.9%) were prominent. Notably, the expression of molecular markers identified within our experiment was consistent with a previous DRN single-cell datasets (**Figure S2B**)^32,33^.

We next aimed at defining the transcriptional signatures of DRN neurons according to their connectivity pattern with the LHA (**Figure 2D, 2E and S2C**). First, our analysis shows that neurons receiving LHA inputs (eGFP+) are predominately GABAergic (51.5%), with smaller proportions of glutamatergic (23.8%) and serotonergic cells (20.8%). In contrast, neurons projecting to the LHA (mCherry+) displayed a more balanced distribution between inhibitory (39.1%) and excitatory (34.3%) neurons, with a minor dopaminergic component (6.0%). Finally, double-labeled DRN neurons included a higher fraction of serotonergic cells (31.7%), consistent with the connectivity patterns described in the previous section.

To describe the gene signatures of eGFP and mCherry expressing DRN neurons, we performed differential expression analysis combined with gene ontology. Comparative transcriptomic analysis identified a set of differently expressed genes (DEGs; FDR < 0.05) distinguishing mCherry+ from eGFP+ neurons, including Hdac5, Cdk8 and Kcnab3, genes associated with neuronal maturation and synaptic vesicle organization (**Figure 2F and 2G**). Hierarchical clustering of DEGs further supported the segregation between connectivity-defined populations. Double-labeled neurons clustered with eGFP+ neurons, whereas mCherry were transcriptionally closer to unlabeled DRN neurons (**Figure 2H**). These findings indicate the DRN neurons receiving LHA inputs form a transcriptionally distinct population, while DRN→LHA neurons more closely resemble the broader neuronal ensemble.

Serotonin neurons in the DRN are highly heterogenous^32^. Thus, to further resolve this heterogeneity, we re-clustered serotonergic cells (sert+) independently. This analysis revealed at least three transcriptionally distinct serotonergic subpopulations (**Figure S3A-S3C**) characterized by selective expression of Zeb2, Oprk1 and Met, genes previously associated with developmentally and functionally diverse 5-HT neuron subtypes^32,33^. Consistent with our anatomical observations (**Figure 1**), serotonergic neurons represented only a small fraction of transgene-positive DRN neurons, indicating that LHA connectivity is primarily established with non-serotonergic populations.

Together, these results show that DRN neurons receiving LHA input constitute a transcriptionally distinct population, predominantly non-serotonergic, and distinguishable from DRN neurons projecting to the LHA.

### LHA provides mixed excitatory and inhibitory monosynaptic inputs to DRN neurons

The LHA sends both excitatory and inhibitory projections to the DRN^12,27^. To quantify the relative strength of these inputs onto serotonergic and non-serotonergic DRN neurons, we expressed ChR2-eYFP in LHA neurons (AAV-ChR2-eYFP) and selectively labeled serotonergic neurons in ePet-cre mice with tdTomato (AAV-DIO-tdTomato in DRN; **Figure 3A and 3B**). Whole-cell voltage-clamp recordings were obtained from tdTomato+ (serotonergic) and tdTomato- (non-serotonergic) DRN neurons in acute slices. Biocytin was included in the internal solution for post hoc identification and immunostaining against TPH (**Figure 3C, 3H and S4).** Monosynaptic transmission was isolated using tetrodotoxin (TTX) and 4-aminopyridine (4-AP). Optical stimulation of LHA terminals reliably evoked postsynaptic currents, recorded at −60mV and 0mV to isolate excitatory (EPSC) and inhibitory (IPSC) postsynaptic current respectively, and pharmacologically confirmed using NBQX and gabazine (**Figure 3E-F and 3J-K**).

**Figure 3.**
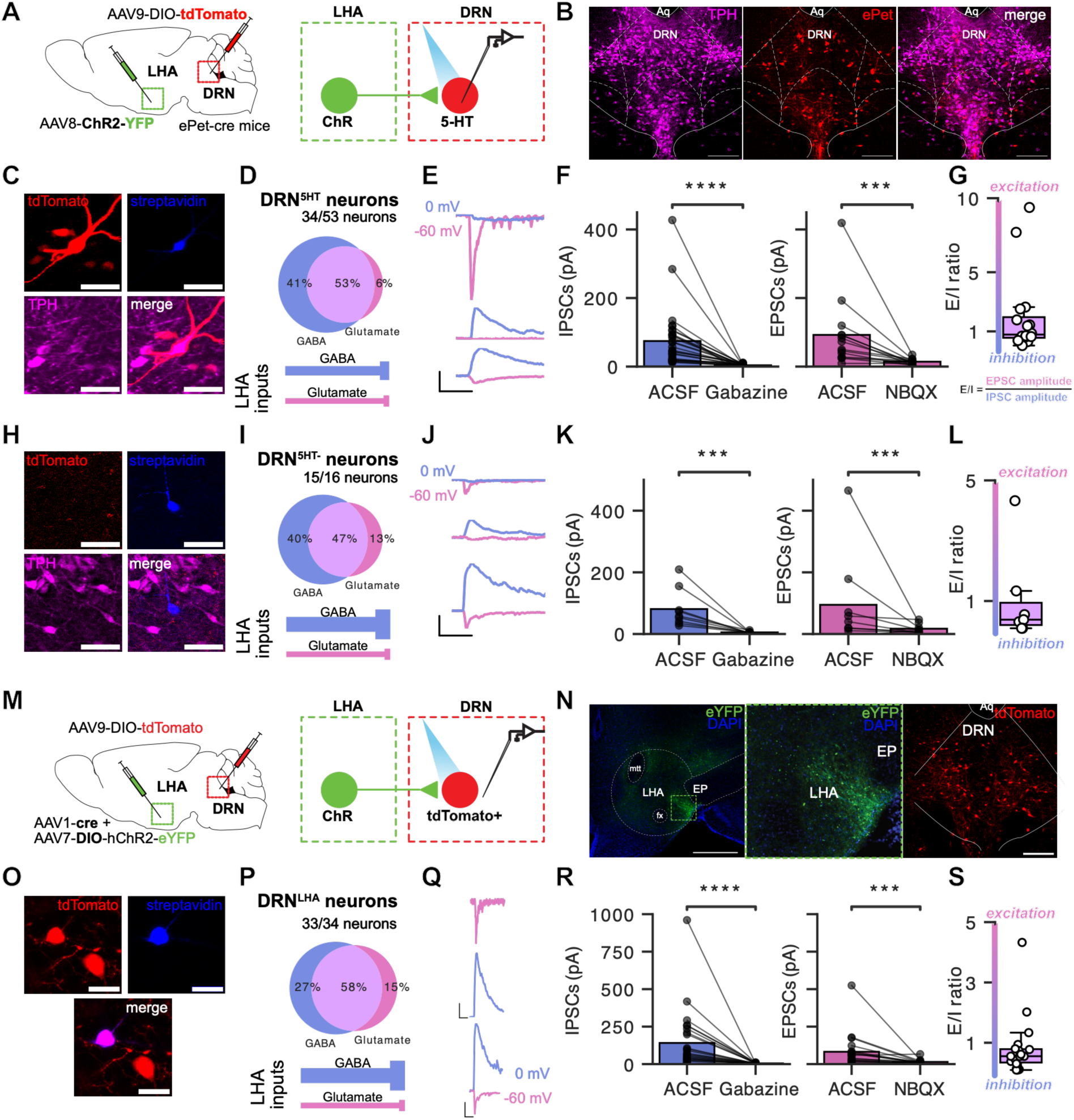
LHA provides mixed excitatory and inhibitory monosynaptic inputs to DRN neurons. A. Experimental design in ePet-Cre mice. AAV-ChR2-YFP was injected in the LHA to express ChR2 in LHA neurons. AAV-DIO-tdTomato was injected in the DRN to label serotonergic (5-HT; TPH+) neurons. Whole-cell recordings were obtained from tdTomato+ (5-HT) and tdTomato− (non-5-HT) neurons during optogenetic stimulation of LHA terminals. B. Representative coronal sections showing tdTomato expression in DRN 5-HT neurons, and co-labeling with TPH immonostaining. Scale bars: 200 µm. C, H. Representative images of recorded tdTomato+ (5-HT; TPH+) and tdTomato- (non-5-HT; TPH−) neurons, respectively. Scale bars: 25 µm. D, I. Proportion DRN 5-HT (D) and non-5-HT (I) neurons exhibiting inhibitory, excitatory or mixed postsynaptic responses following LHA terminals stimulation. E, J. Representative voltage-clamp traces recorded at −60 mV (EPSCs) and 0 mV (IPSCs) in 5-HT (E) and non-5-HT (J) neurons following LHA terminals stimulation, F, K. Amplitudes of optogenetically-evoked IPSCs and EPSCs recorded in ACSF and following bath application of gabazine (10 µM) or NBQX (10 µM) in 5-HT (F) and non-5-HT (K) neurons. GABAzine abolished IPSCs, and NBQX abolished EPSCs. G, L. Distribution of excitation/inhibition (E/I) ratios in neurons exhibiting both IPSCs and EPSCs in 5-HT (G) and non-5-HT (L) populations. M. Experimental design in wild-type mice to selectively recorded from DRN neurons receiving LHA inputs (DRN^:LHA^). ChR2 was expressed in LHA neurons and tdTomato was expressed in DRN neurons receiving LHA inputs. N. Representative images of ChR2 expression in LHA and tdTomato labeling in DRN receiving inputs from LHA. Scale bars: 200 µm. O. Representative image of a patched DRN^:LHA^ neuron. Scale bar: 20 µm. P. Proportion of inhibitory, excitatory, and mixed postsynaptic responses in DRN^:LHA^ following LHA terminal stimulation. Q. Representative voltage-clamp traces from DRN^:LHA^ recorded at −60 mV and 0 mV following LHA terminal stimulation. R. Amplitudes of optogenetically evoked IPSCs and EPSCs in ACSF and following bath application of gabazine (10 µM) or NBQX (10 µM). S. Distribution of E/I ratios in DRN^:LHA^ neurons exhibiting both IPSCs and EPSCs. Statistical comparisons were performed using paired tests for pharmacological blockage and appropriate non-parametric or parametric tests as described in Methods. \*\*\**p* < 0.001 and \*\*\*\**p* < 0.0001.

Among serotonergic neurons (tdTomato+), 64% (34 out of 53) exhibited monosynaptic responses to optical stimulation of LHA terminals. Of these, 41% received purely inhibitory input, 6% purely excitatory input, and 53% mixed excitatory and inhibitory input (**Figure 3D**). For neurons receiving mixed inputs, the median excitation-to-inhibition (E/I) ratio was 0.8, indicating a relative predominance of inhibitory synaptic drive under these recording conditions (**Figure 3G**).

Non-serotonergic neurons (tdTomato-) responded more frequently (94%; 15/16), with 40% receiving purely inhibitory input, 13% purely excitatory input, and 47% mixed input (**Figure 3I)**. These neurons exhibited a stronger inhibitory bias, with a median E/I ratio of 0.4 (**Figure 3L**). Together, these results demonstrate that LHA provides direct monosynaptic input to both serotonergic and non-serotonergic DRN neurons, with inhibitory transmission predominating, particularly in non-serotonergic populations.

To validate the AAV1-cre tracing strategy (used in **Figures 1 and 2**), we selectively expressed ChR2 in LHA neurons and labeled their postsynaptic DRN targets with tdTomato using AAV1-cre-dependent recombination (**Figure 3M-N**). Optical stimulation of LHA terminals evoked monosynaptic responses in 97% of tdTomato+ neurons (33/34 cells; **Figure 3P-S**), with a distribution of inhibitory, excitatory, and mixed responses comparable to that observed in randomly recorded DRN neurons (**Figure 3P**). These results confirm that AAV1-cre reliably labels DRN neurons receiving functional monosynaptic input from the LHA without strong bias toward specific cell types, validating its use in our anatomical and transcriptional mapping experiments.

### LHA-innervated DRN neurons provide strong local inhibition within the DRN

To determine whether DRN neurons innervated by the LHA (DRN^:LHA^) participate in local DRN microcircuits, we expressed ChR2 selectively in DRN^:LHA^ using AAV1-Cre in the LHA and AAV-DIO-ChR2-eYFP in the DRN (**Figure 4A**). Whole-cell voltage-clamp recordings were obtained from non-eYFP DRN neurons while optically stimulating ChR2-expressing DRN^:LHA^ neurons in the presence of TTX and 4-AP.

**Figure 4.**
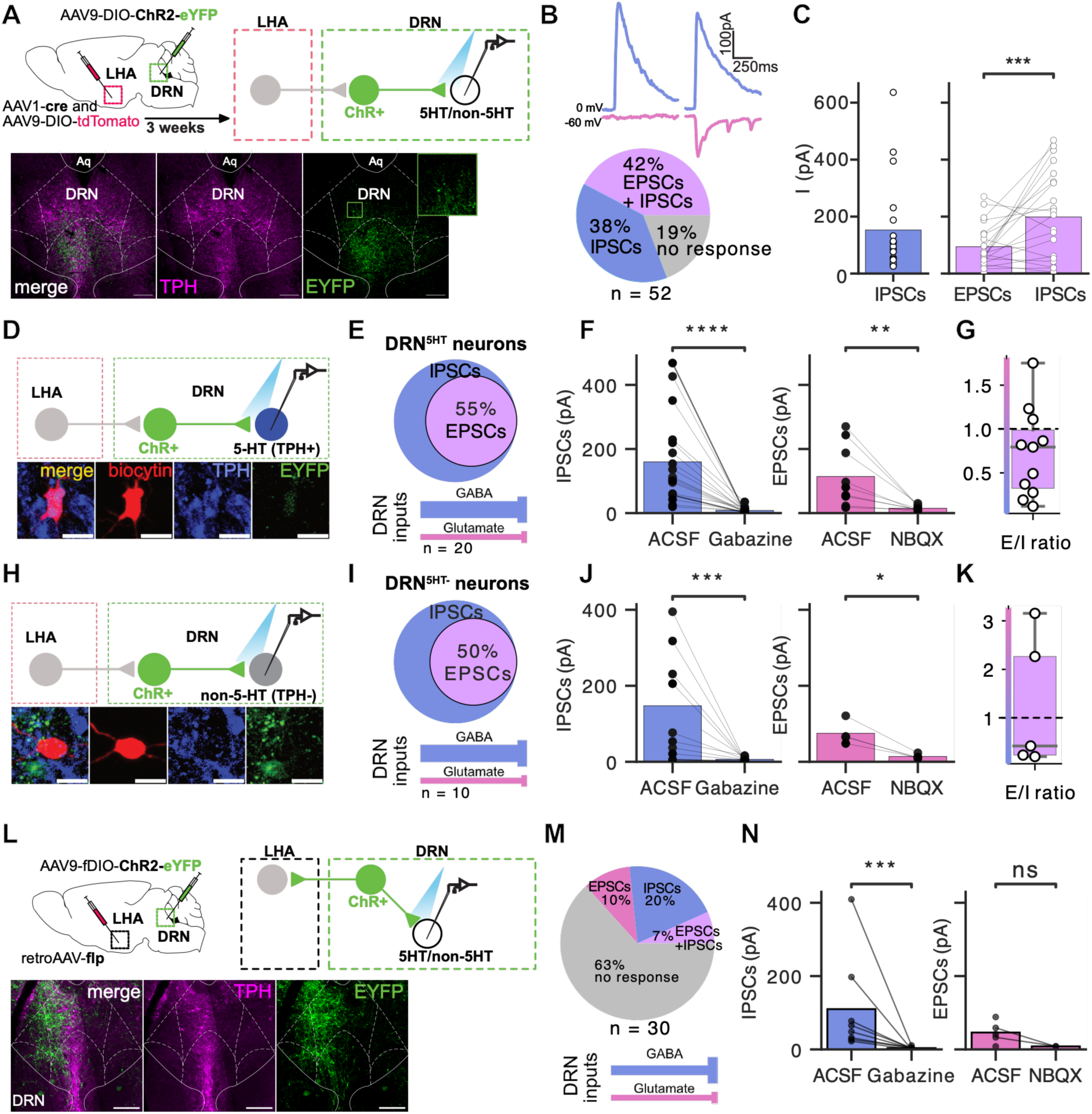
LHA-innervated DRN neurons provide strong local inhibition within the DRN. A. Experimental strategy to assess intra-DRN connectivity of LHA-innervated neurons (DRN^:LHA^). ChR2-eYFP was expressed selectively DRN neurons receiving LHA inputs. Whole-cell recordings were obtained from randomly sampled DRN neurons, followed by post hoc TPH immunostaining to identify serotonergic (5-HT) and non-serotonergic (5-HT-) neurons. Scale bars: 200 µm (overview). B. Representative coltage-clamp traces recorded at −60 mV and 0 mV in response to optogenetic stimulation of DRN^:LHA^ neurons, and proportion of responsive cells (n = 52). C. Distribution of optogenetically evoked EPSCs and IPSCs amplitudes recorded in ACSF in the presence of 1 µM TTX and 100 µM 4-AP, confirming monosynaptic connectivity. D, H. Representative images of recorded TPH+ (D) and TPH− (H) neurons. Scale bar: 25 µm. E, I. Proportion of TPH+ (E) and TPH- (I) neurons exhibiting inhibitory or mixed postsynaptic responses following stimulation of DRN^:LHA^ neurons. F, J. Amplitudes of optogenetically evoked IPSCs and EPSCs in ACSF and following bath application of gabazine (10 µM) or NBQX (10 µM) in TPH= (F) and TPH- (J) neurons. G, K. Distribution of E/I ratios in neurons exhibiting both IPSCs and EPSCs. L. Experimental strategy to assess intra-DRN connectivity of DRN neurons projecting to the LHA (DRN→LHA). ChR2 was expressed selectively in DRN→LHA neurons, and whole-cell recordings were obtained from randomly sampled DRN neurons. Scale bars: 200 µm. M. Proportion of postsynaptic responses evoked by optogenetic stimulation of DRN→LHA neurons. N. Distribution of evoked IPSC and EPSCs amplitude recorded in ACSF in the presence of 1 µM TTX and 100 µM 4-AP following stimulation of DRN→LHA neurons. Statistical comparisons were performed using paired tests for pharmacological blockage and appropriate non-parametric or parametric tests as described in Methods. \**p* < 0.05, \*\**p* < 0.01, \*\*\**p* < 0.001 and \*\*\*\**p* < 0.0001.

Of 52 recorded neurons, 42 (∼80%) exhibited postsynaptic responses, indicating extensive intra-DRN connectivity (**Figure 4B**). Among the responsive neurons, all exhibited inhibitory postsynaptic currents, and 52% (22/42) also displayed concurrent excitatory currents of smaller amplitude (**Figure 4B-C**).

Both serotonergic (TPH+, n = 20) and non-serotonergic (TPH-, n = 10) neurons received inhibitory inputs, with approximately half of each group also receiving mixed excitation and inhibition (**Figure 4E-4I and S5**). For mixed-inputs cells, median E/I ratios were 0.8 for serotonergic neurons and 0.4 for non-serotonergic neurons, indicating a predominant inhibitory influence **(Figure 4G and 4K).**

In contrast, DRN neurons projecting to the LHA (DRN→LHA) exhibited substantially fewer local connections. Using our intersectional viral approach to selectively express ChR2 in DRN→LHA neurons, we found that only 37% of recorded neurons (11/30) displayed any postsynaptic response, and the majority showed no detectable local connectivity (63%; 19/30). Among the responsive neurons, 27% (3/11) exhibited excitatory postsynaptic current, 55% (6/11) exhibited inhibitory current and 18% (2/11) displayed concurrent excitatory and inhibitory currents **(Figure 4l-N)**. This marked difference indicates that DRN neurons receiving LHA input are more strongly embedded within local DRN circuitry, whereas DRN→LHA neurons primarily function as long-range projection neurons with limited local integration.

### Chronic silencing of LHA-targeted DRN neurons induces hyperlocomotion and repetitive behaviors

To assess the behavioral contribution of DRN^:LHA^ neurons, we chronically silenced their synaptic output by expressing tetanus toxin light chain (TeLC) selectively in these neurons (**Figure 5A**). TeLC cleaves the vesicle-associated protein synaptobrevin, thereby blocking neurotransmitter release^34^. Control mice expressed eYFP alone. After 6 weeks of expression, mice underwent a behavioral battery assessing locomotion, stress-related coping, anxiety-like behavior, social interaction, repetitive behavior, and consummatory behavior (**Figure 5B**).

**Figure 5.**
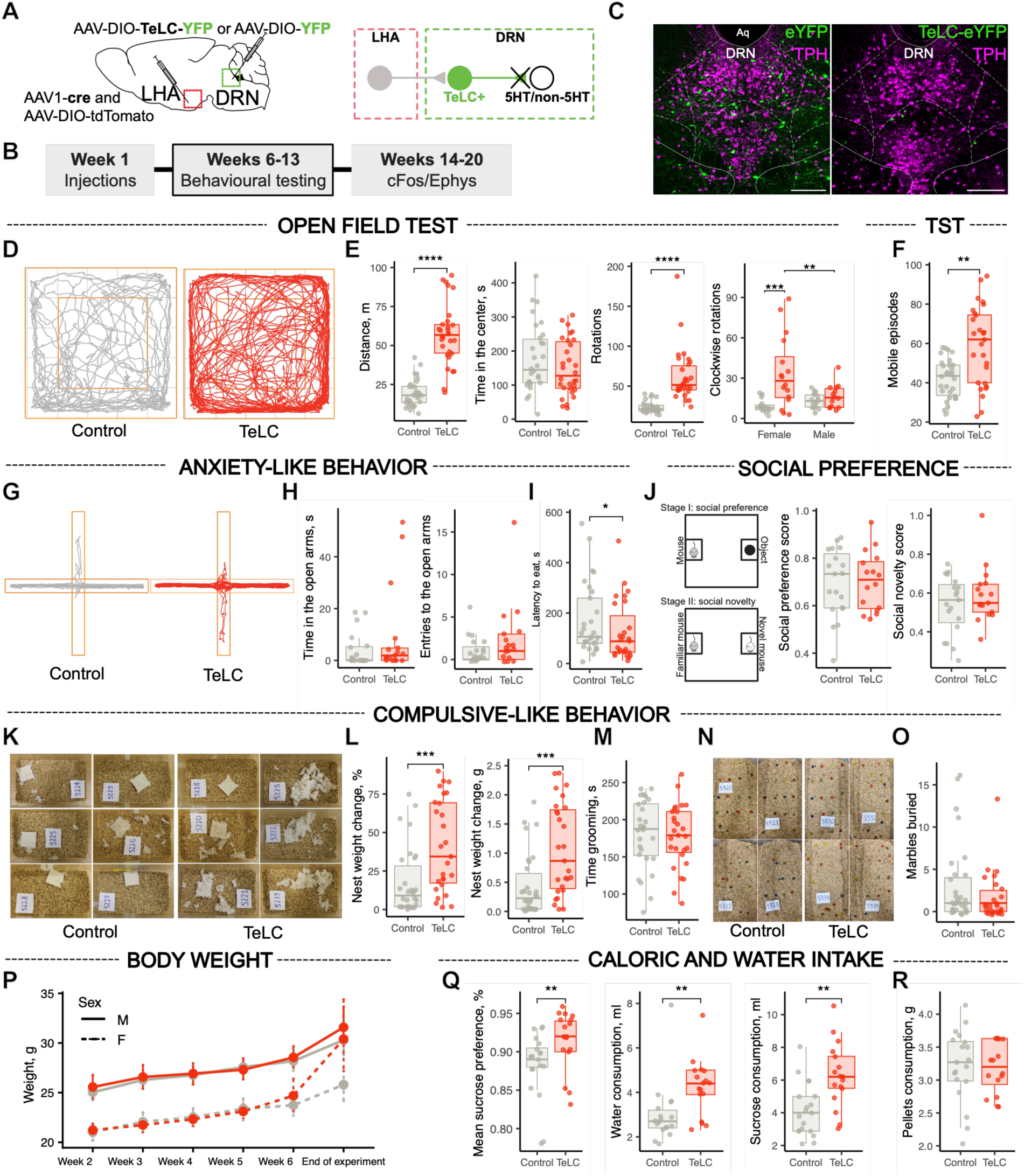
Chronic silencing of LHA-targeted DRN neurons induces hyperlocomotion and repetitive behaviors. A. Schematic of experimental strategy. DRN neurons receiving LHA input were chronically silenced by Cre-dependent expression of tetanus toxin light chain (TeLC-eYFP); control mice expressed eYFP alone. B. Experimental timeline. Behavioral testing was conducted six weeks after viral injections, followed by electrophysiology and cFos mapping. C. Representative confocal images of DRN neurons expressing YFP (controls) or TeLC-YFP. TPH+ immunostaining is shown in magenta. Scale bar: 200 µm. D. Representative open field test (OFT) locomotor tracks from control and TeLC mice. E. OFT quantification: total distance travelled, time spent in the center zone, total number of rotations, and rotation direction analyzed by sex (n= 60). F. Tail suspension test (TST): number of mobile episodes (n= 57). G. Representative elevated plus maze (EPM) tracks. H. EPM quantification: time spent in the open arms and number of open arm entries (n= 35). I. Novelty-suppressed feeding test (NSF): latency to initiate eating (n = 56). J. Social preference and social novelty scores (n= 35). K. Representative nests following the nest shredding assay. L. Nest shredding: change in nest weight after 1 h (n = 57). M. Splash test: time spent grooming (n = 55). N. Representative marble burying test images. O. Marble burying: number of marbles buried (n = 57). P. Body weight measured weekly following viral injection (n = 56). Q. Sucrose preference and fluid consimption (n =36). R. Pelleted food consumption (n = 35). Boxplots show median and interquartile range (IQR), with whiskers extending to 1.5× IQR; individual points represent mice. Line plots show the mean ± bootstrapped 95% confidence intervals. Statisitcal analyses were performed using linear models, negative binomial GLMs with log link, or Gamma GLMs with log link as appropriate, including treatment, sex, and their interaction, adjusted for batch (see details in Supplementary tables for full statistics). Significance levels are indicated in panels. \**p* < 0.05, \*\**p* < 0.01, \*\*\**p* < 0.001 and \*\*\*\**p* < 0.0001.

In the open-field test (OFT), TeLC mice displayed robust hyperlocomotion compared to controls, characterized by increased total distance traveled, higher mean movement speed, and reduced immobility episodes (**Figure 5D-E** and **S6A**). This behavioral activation was accompanied by a marked reorganization of movement structure: TeLC mice exhibited a reduced number of discrete movement bouts, driven by a significant increase in average bout duration, indicating more continuous and sustained locomotion rather than frequent start-stop movement. TeLC mice also displayed pronounced circling behavior, observed in both sexes, with sex-dependent rotation direction (**Figure 5D-E and 6SA**). With respect to spatial exploration, TeLC mice made more frequent entries into the center zone of the arena but spent less time per visit in this region (**Figure S6A**). Consequently, total time spent in the center zone did not differ significantly from controls, indicating that increased locomotor activity was not associated with altered anxiety-related exploration. Together, these measures indicate that TeLC mice engage in persistent, high-speed locomotion with reduced pausing, consistent with a state of heightened motor drive.

In the tail-suspension test (TST), TeLC exhibited increased mobility, reflected by a higher number of active episodes and elevated mobility score (**Figure 5F and S6B**), indicating enhanced spontaneous motor output under stress.

Because repetitive circling behavior has been described in mouse models of anxiety, obsessive-compulsive disorder (OCD) and autism^35,36^, we next examined whether the hyperlocomotor phenotype observed in TeLC mice was accompanied by broader alterations in anxiety-like, social, or repetitive behaviors.

TeLC mice did not differ from controls in anxiety-like behavior in the elevated plus maze, nor did they show alterations in social preference, social novelty, or direct social interaction (**Figure 5G-J and Figure S7**). However, TeLC showed a significant decrease in latency to eat in the novelty-suppressed feeding test (**Figure 5I)**. These findings indicate that chronic silencing of DRN^:LHA^ neurons selectively affects locomotor and activity-related behaviors without producing generalized anxiety or social deficits.

Assessment of repetitive behaviors revealed increased nestlet shredding in TeLC mice, whereas grooming behavior and marble-burying were unchanged (**Figure 5K-O**). Together with circling behavior, these findings indicate a bias toward repetitive motor actions. However, given the absence of changes in other compulsivity-associated assays, these behaviors are more conservatively interpreted as hyperlocomotor and repetitive rather than compulsive per se.

Body weight remained comparable between groups, although a subset of females TeLC mice exhibited late-phase weight gain (**Figure 5P**). In the sucrose-preference test, TeLC mice showed increased sucrose preference and consumed greater volumes of sucrose and water solutions, without differences in solid food intake (**Figure 5Q-R**), suggesting increased liquid consumption and sucrose preference without generalized hyperphagia.

### Silencing of LHA-targeted DRN neurons is associated with increased serotonergic activity and reduced local inhibition

To investigate how chronic silencing of the DRN^:LHA^ neurons impacts local DRN activity, we performed cFos mapping and *ex vivo* slice electrophysiology following completion of behavioral testing. Two complementary experimental approaches were used in the same cohort of mice: one assessed spontaneous synaptic transmission in acute slices, and the other quantified neuronal activation under basal (home-cage) conditions and following acute stress induced by TST (**Figure 6A and 6B**).

**Figure 6.**
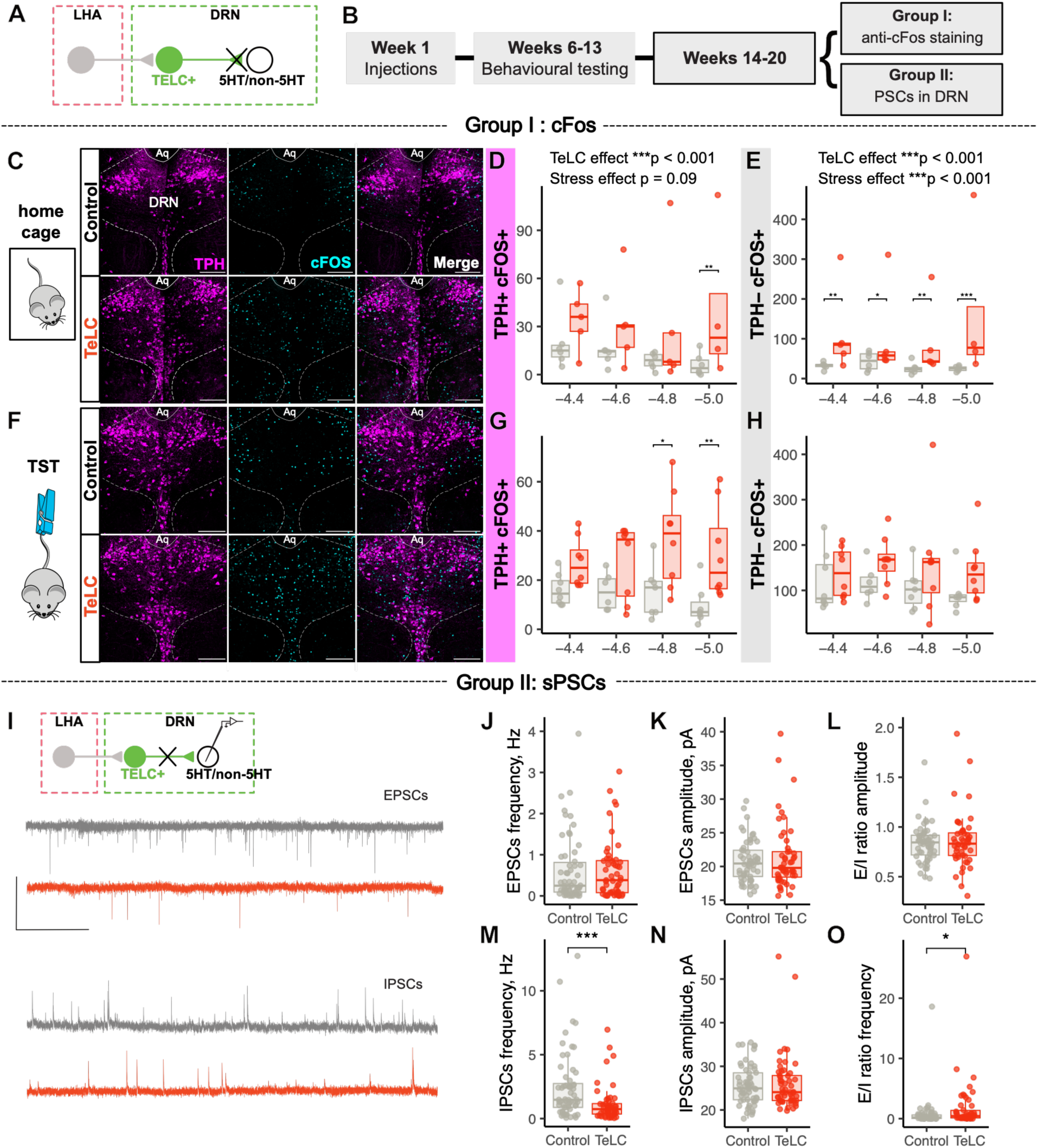
Silencing of LHA-targeted DRN neurons is associated with increased serotonergic activity and reduced local inhibition. A. Schematic of experimental strategy. DRN neurons receiving LHA input were chronically silenced by Cre-dependent expression of tetanus toxin light chain (TeLC-eYFP); control mice expressed eYFP alone. B. Experimental timeline. Following behavioral testing, mice were divided into two groups: Group I for cFos immunostaining and Group II for electrophysiological recordings. C, F. Representative confocal images of DRN coronal sections showing cFos (cyan) and TPH (magenta) immunoreactivity in control (YFP) and TeLC mice under home cage condition (C) or following acute stress (tail suspension test, TST; F). Scale bar: 200 µm. D–E. Quantification of cFos⁺ DRN neurons across the anterior–posterior axis under home cage conditions for TPH⁺ (serotonergic) and TPH⁻ (non-serotonergic) populations. The distance from bregma (mm) is indicated on the x-axis. G–H. Quantification of cFos⁺ DRN neurons across the anterior–posterior axis following TST for TPH⁺ and TPH⁻ populations. The distance from bregma (mm) is indicated on the x-axis. I. Experimental approach for acute slice recordings. Representative traces of spontaneous EPSCs and IPSCs recorded from DRN neurons. J-K. Frequency and amplitude of spontaneous EPSCs. M-N. Frequency and amplitude of spontaneous IPSCs. L, O. Excitation/Inhibition ratios calculated from sPSC amplitudes (L) and frequencies (O). Boxplots show the median and interquartile range (IQR), with whiskers extending to 1.5× IQR; individual points represent mice. Line plots show the mean ± bootstrapped 95% confidence intervals. Statisitcal analyses were performed using linear models, negative binomial GLMs with log link, or Gamma GLMs with log link as appropriate, including treatment, sex, and their interaction, adjusted for batch (see details in Supplementary tables for full statistics). Significance levels are indicated in panels. \**p* < 0.05, \*\**p* < 0.01, and \*\*\**p* < 0.001.

Across the rostrocaudal extent of the DRN, TeLC expression produced a marked increase in neuronal activation as measured by cFos expression. When averaged across all sample sections and conditions (home cage vs TST), TeLC mice exhibited a significant increase in cFos expression in serotonergic (TPH+) neurons (estimated 4.5-fold increase) and non-serotonergic (TPH-) neurons (estimated 3.8-fold increase) compared with controls, whereas acute stress produced a significant 3.7-fold increase only in TPH-population (Ctrl and TeLC groups), no significant change was measured in TPH+ neurons (**Figure S8**).

In serotonergic neurons, pairwise comparisons revealed that TeLC-associated increases in cFos expression were not uniform along the rostrocaudal axis. Under basal conditions, increased cFos labeling was observed primarily in the most caudal DRN (−5.0 mm from bregma). Following TST, elevated cFos expression in TeLC mice extended to both mid-caudal and caudal levels (−4.8 and −5.0 mm from bregma), with a significant interaction between TeLC expression and stress at −4.8 **(Figure 6D and 6G)**. Although interaction effects were not detected at all rostrocaudal levels, these data indicate that DRN^:LHA^ silencing alters the spatial pattern of serotonergic activation, preferentially engaging caudal DRN subregions (**Figure S8D**).

In non-serotonergic neurons, TeLC expression increased cFos labeling across all rostrocaudal levels under basal conditions. In contrast, following TST, cFos expression in TPH- neurons was similarly elevated in both control and TeLC mice (Figure 6E and 6H), indicating that non-serotonergic DRN neurons are strongly stress-reactive and that stress occludes the effect of DRN^:LHA^ silencing in this population. We detected significant TeLC x stress interactions at −4.6 and −4.8 mm (**Figure S8E)**.

Together, these findings demonstrate that chronic silencing of DRN^:LHA^ neurons elevates baseline activity of both serotonergic and non-serotonergic DRN populations, with serotonergic activation exhibiting pronounced regional specificity within the mid-caudal and caudal DRN.

To assess whether DRN^:LHA^ silencing was associated with changes in local synaptic transmission, we recorded spontaneous excitatory and inhibitory postsynaptic current (sEPSCs abd sIPSCs) from DRN neurons in acute slices (**Figure 6I-O**). Compared with controls, TeLC mice exhibited a significant reduction in sIPSCs frequency, while sIPSC amplitude and both sEPSC frequency and amplitude were unchanged. This reduction was observed in both TPH- and TPH+ (**Figure S8F-H**). Consequently, the excitation-to-inhibition (E/I) frequency ratio was increased, reflecting a reduction in inhibitory tone within the DRN. These results indicate that chronic TeLC expression in DRN^:LHA^ neurons decreases inhibitory input onto neighboring DRN neurons, consistent with a disinhibitory mechanism that may contribute to the elevated neuronal activity observed *in vivo*.

Overall, chronic silencing of DRN^:LHA^ neurons was associated with reduced local inhibitory synaptic tone and elevated activity in the DRN, notably for serotonergic populations within the mid-caudal regions, supporting a disinhibitory mechanism linking LHA input to serotonergic regulation.

## DISCUSSION

The LHA exerts broad influence over behavioral state through its projections to multiple neuromodulatory centers, yet how hypothalamic signals interface with the heterogeneous circuitry of the DRN has remained unclear. Here, we delineate a circuit organization through which the LHA can influence serotonergic activity and behavioral activation via a local inhibitory microcircuit within the DRN. Using a combination of connectivity-based viral tracing, electrophysiology, snRNA sequencing, and behavioral manipulations, we show that LHA inputs preferentially engage non-serotonergic DRN neurons that are strongly embedded in local inhibitory networks. Chronic silencing of these LHA-innervated DRN neurons was associated with reduced local inhibitory synaptic tone, elevated activity of serotonergic populations, particularly in caudal DRN subregions, and increased locomotor output in neutral and stress contexts. Together, these findings support a model in which hypothalamic input biases serotonergic function indirectly, through suppression of intra-DRN inhibition, rather than through direct excitation of 5-HT neurons.

### LHA inputs preferentially target non-serotonergic and transcriptionally distinct DRN populations

The DRN receives dense hypothalamic innervation, including both glutamatergic and GABAergic projections from the LHA^12,27^, but the cellular organization and functional logic of these inputs have been poorly defined. By combining anterograde and retrograde viral tracing with molecular profiling, we found that LHA axons predominantly innervate non-serotonergic DRN neurons. Across anatomical and transcriptomic analysis, serotonergic neurons accounted for only a minority of LHA-innervated DRN cells, whereas the majority expressed markers of inhibitory neurotransmission, with smaller glutamatergic and minor dopaminergic subsets.

Importantly, snRNA sequencing revealed that DRN neurons defined by their connectivity with the LHA are not simply a random sampling of inhibitory neurons. Instead, LHA-innervated DRN neurons occupied a transcriptionally distinct molecular space relative to both DRN neurons projecting back to the LHA and the broader DRN neuronal population. This suggest that LHA inputs selectively engage a functionally specialized inhibitory ensemble rather than generic GABAergic neurons. Such connectivity-defined molecular segregation supports the idea that the long-range afferents access the DRN through dedicated inhibitory microcircuits, adding a layer of functional organization beyond classical transmitter identity.

This organization parallel that reported for other major DRN afferents, including projections from the medial prefrontal cortex, the lateral habenula, the preoptic area and the Edinger-Westphal nucleus, which also preferentially target non-serotonergic DRN populations to influence serotonergic outputs^13,37–39^. Together, these findings reinforce a view of DRN GABAergic neurons as integrative nodes through which diverse forebrain and brainstem inputs modulate serotonergic tone in a context-dependent manner.

### Synaptic organization supports indirect modulation of serotonergic neurons

Having established that LHA inputs preferentially innervate non-serotonergic neurons, we next examined how this connectivity is reflected at the level of synaptic transmission. Optical stimulation of LHA terminals evoked monosynaptic responses in the majority of non-serotonergic DRN neurons and in a substantial fraction of serotonergic neurons, with inhibitory transmission predominating in both populations. Although mixed excitatory and inhibitory inputs were observed in a subset of cells, the overall synaptic balance was biased toward inhibition. At the network level, this configuration is consistent with a circuit in which activation of LHA inputs preferentially suppress inhibitory DRN neurons, thereby reducing their influence over serotonergic populations.

Consistent with this idea, mapping of Click or tap here to enter text.intra-DRN connectivity revealed that LHA-innervated DRN neurons form extensive local inhibitory connections. Approximately 80% of recorded DRN neurons exhibited postsynaptic responses to stimulation of this population. All responsive neuron received inhibitory inputs, and approximately half also displayed concurrent excitatory current, with inhibition predominating in mixed-response cells. These findings indicate that the LHA-innervated DRN neurons are strongly embedded within local DRN circuitry and constitute a major source of inhibitory control. In contrast, DRN neurons projecting back to the LHA (DRN→LHA) formed comparatively sparse local connections and were transcriptionally closer to the general DRN population, suggesting that they primarily function as long-range projection neurons than local integrators.

Moreover, chronic TeLC-mediated silencing of LHA-innervated DRN neurons reduced the frequency of spontaneous inhibitory synaptic events without altering excitatory transmission, shifted the local excitation-inhibition balance, and increased cFos expression in the DRN, notably in serotonin neurons in the most caudal DRN. Given that LHA inputs to the DRN are predominantly inhibitory and preferentially engage non-serotonergic neurons embedded in local inhibitory circuits, these findings provide convergent support for a disinhibitory mechanism within the DRN. In this framework, LHA-innervated inhibitory neurons normally constrain serotonergic activity, and their disruption relieves this local inhibition, thereby biasing DRN output toward enhanced serotonergic activation.

### Reciprocal DRN-LHA connectivity and potential bidirectional control

Our data further indicate that reciprocal DRN-LHA connectivity is organized through largely distinct neuronal populations. DRN neurons projecting to the LHA showed minimal overlap (∼10%) with LHA-innervated DRN neurons, differed in their spatial distribution within the DRN, and exhibited limited local connectivity. This segregation suggests a functional asymmetry between input-defined and output-defined DRN populations, with LHA-innervated neurons positioned to shape local circuit dynamics and DRN→LHA neurons serving primarily as long-range signaling elements.

Previous studies have shown that DRN GABAergic projections to the LHA or VTA promote wakefulness^40,41^, whereas projections to the paraventricular thalamus reduce arousal^42^. In this context, one possibility is that reciprocal DRN-LHA loops coordinate behavioral states through complementary inhibitory and excitatory pathways. LHA inputs may suppress local inhibitory constraints within the DRN, while DRN outputs to the LHA could feed back information about internal state or ongoing behavior. Although this hypothesis remains to be tested, it highlights the potential for bidirectional interactions between hypothalamic and raphe circuits in regulating arousal and behavioral activation.

### Behavioral consequences of disrupting LHA-innervated DRN circuitry

Behaviorally, chronic silencing of LHA-innervated DRN neurons produced robust hyperlocomotion and increased mobility in neutral and stress contexts, without affecting anxiety-like or social behaviors. This pattern is consistent with previous reports showing that chronic activation of DRN serotonergic neurons, as well as by pharmacological enhancement of serotonin transmission, increases locomotor output and mobility in aversive contexts such as in forced- or tail-suspension tests^8–10,43^. In our experiments, these behavioral changes were accompanied by increased cFos expression in serotonergic neurons, particularly within mid-caudal and caudal DRN subregions, suggesting an association between disruption of local inhibitory circuitry and elevated serotonergic activity.

Importantly, similar increases in locomotor activity have been observed following chemogenetic or optogenetic inhibition, as well as genetic ablation of DRN GABAergic neurons^42,44^, indicating that local inhibitory tone within the DRN normally constrains behavioral activation. Consistent with this idea, optogenetic inhibition of DRN GABA neurons increases movement without altering anxiety-related behavior in the open field, closely paralleling the phenotype observed following TeLC-mediated silencing of LHA-innervated DRN neurons^44^. Together, these findings suggest that disruption of inhibitory control within DRN microcircuits selectively biases behavioral output toward heightened motor activation.

Previous work has also linked DRN GABA neurons to arousal regulation, showing that stress activates these neurons^13,37^ and that their ablation increases both locomotion and arousal (as measured by pupil dilation)^42^. In this context, LHA inputs may participate in regulating arousal-related behavioral states by modulating DRN inhibitory circuit and, indirectly serotonergic tone during stressClick or tap here to enter text.. Such a mechanism could account for the combination of increased locomotion, stress-related mobility, circling behavior, and increased nest-shredding observed here, consistent with enhanced behavioral activation and repetitive motor output, with features reminiscent of compulsive-like behaviors but without anxiety-related behavior.

Notably, the regional specificity of serotonergic activation observed in the caudal DRN is consistent with prior work demonstrating functional heterogeneity along the rostro-caudal axis of the DRN, with caudal subregions preferentially implicated in locomotor and arousal-related processes^1,32^. The selective engagement of these serotonergic populations suggest that LHA-driven disinhibitory mechanisms may bias specific DRN output channels involved in behavior control.

### Integration with hypothalamic disinhibitory motifs

The disinhibitory architecture identified in the LHA→DRN pathway mirrors circuit motifs described for other hypothalamic outputs. LHA GABAergic projections to the VTA disinhibit dopamine neurons through suppression of local GABA cells, promoting reinforcement and behavior activation^22,24^. Notably, a substantial fraction of DRN-projecting LHA neurons collateralizes to the VTA, and coordinated activity in these pathways coincides with increased dopaminergic and serotonergic activity at movement initiation^27^. These observations support a broader principle in which hypothalamic inhibitory outputs bias behavioral state by engaging local disinhibitory microcircuits within downstream neuromodulatory centers. Within this framework, the LHA→DRN circuit described here likely operates in parallel with the LHA→VTA pathway to coordinate serotonergic and dopaminergic contributions to behavioral activation. Rather than broadcasting a unitary motivational signal, hypothalamic inhibitory projections may provide a state-dependent activation bias, flexibly engaging distinct neuromodulatory systems depending on context and behavioral demands.

### Limitations and future directions

Several limitations should be considered. First, although AAV1-Cre primarily labels postsynaptic targets, limited retrograde spread cannot be fully excluded^28^. However, the high proportion of optogenetically responsive DRN neurons and anatomical segregation of labeling argue against substantial contamination. Second, TeLC silencing affect all synaptic outputs of targeted neurons, precluding definitive separation of local versus long-range contributions to the observed behavioral effects. Third, while relatively few, neurons infected outside the DRN could not be entirely excluded and may have contributed modestly to the observed effects. Future work will be needed to determine how excitatory LHA inputs to the DRN contribute to behavior. These projections likely complement the inhibitory LHA pathway described here, supporting distinct aspects of behavioral control. In previous work, we found that activity of LHA terminals in the DRN signals aversive cues and their predictors during an active avoidance task, consistent with a role in associative learning^27^. Thus, glutamatergic LHA→DRN projections may contribute to aversive learning and the orchestration of defensive behaviors, similar to the established roles of glutamatergic outputs to the VTA^24–26^. Together with GABAergic LHA projections, these complementary pathways may help balance defensive and activating behavioral states.

## Conclusion

In summary, our findings support a model in which the LHA influences serotonergic function through engagement of a local inhibitory microcircuit within the DRN. By preferentially targeting transcriptionally distinct non-serotonergic DRN neurons that are strongly embedded in local inhibitory networks, LHA inputs are positioned to bias serotonergic activity indirectly. This circuit organization parallels disinhibitory motifs used by LHA to engage other monoaminergic systems and provide a framework for understanding how hypothalamic signals coordinate behavioral activation through parallel modulation of serotonergic and dopaminergic pathways.

## ACKNOWLEDGMENTS

We thank Dr Silvia Pozzi’s laboratory and Quentin Leboulleux for their technical assistance. We kindly thanks Dr Kunzhang Lin and Dr Fuqiang Xu for providing the plasmids for AAV11. We also thank the CERVO Canadian Optogenetics and Vectorology Foundry Core Facility for producing the viral vectors. C.D.P. was supported by the Canadian Institutes of Health Research grant PJT169117 and is supported by the Natural Science and Engineering Research Council of Canada grant RGPIN-2025-04515, and receives Fonds de Recherche en Santé du Québec (FRQS) Junior-2 salary support. R.S. was supported by a doctoral training scholarship from FRQS.

## AUTHOR CONTRIBUTIONS

R.S. and C.D.P. conceived, supervised and directed the study. R.S., Z.B., V.E., C.Z., and M.P performed the experiments. R.S. and A.M.R. performed the formal analysis of the data. R.S. and C.D.P. interpreted the data. R.S. and C.D.P. wrote and edited the manuscript. All authors provided feedback on the manuscript.

## DECLARATION OF INTEREST

The authors declare no competing interests.

### Declaration of generative AI and AI-assisted technologies

During the preparation of this work, the authors used ChatGPT to check grammar and spelling. After using this tool, the authors reviewed and edited the content as needed. No generative AI tools have been used to produce any new content. The authors take full responsibility for the content of the publication.

## METHODS

### Resource availability

Further information and requests for resources should be directed to and will be fulfilled by the lead contact Christophe Proulx.

### Material availability

This study did not generate new unique reagents.

### Experimental model and study participant details

Both males and females were used in this study (20-30 g). All experiments were performed with 8- to 24-week-old wild-type C57Bl/6 or ePet-cre mice. The mice were housed 2-4 per cage and were kept on a 12 h/12 h light/dark cycle. All the experiments were performed in accordance with the Canadian Guide for the Care and Use of Laboratory Animals guidelines and were approved by the Université Laval Animal Protection Committee.

### Stereotactic injections

The mice were anesthetized with isoflurane for the stereotaxic injections of adeno-associated viruses (AAVs). Viral titers ranged from 10^12^ to 10^13^ genome copies per milliliter and volumes ranged from 100 to 200 nl per side. Injections were performed using a glass pipette mounted on a stereotactic table. The AAVs were infused at a rate of 1 nl/sec. At the end of the injection, the pipet was left *in situ* for 5 min to allow the virus to diffuse into the surrounding tissue.

To label DRN neurons which are synaptically contacted by LHA axons, the mice were injected with a mixture of AAV2/retro-CAG-flpO-P2A-TagBFP2 and AAV1-CAG-cre-WRPE in LHA, and with a mixture of AAV9-DIO-eGFP and AAV9-fDIO-mCherry in DRN. In alternative strategies, instead of AAV2/retro-CAG-flpO-P2A-TagBFP2 we used either CTb647 (Alexa Fluor 647-conjugated cholera toxin subunit B) or AAV2/11-CAG-FlpO-T2A-TagBFP2. The mice were transcardially perfused 3 weeks after the viral injection, and their brains were processed for histology.

For the electrophysiological characterization of LHA-DRN signaling, the mice were injected with a mixture of AAV1-hSyn-cre-WPRE and AAV2/7-EF1a-DIO-hChR2(H134R)-eYFP in LHA (1:1), and with AAV2/9-CAG-Flex-dTomato in DRN.

For the electrophysiological characterization of the local DRN connectivity, the mice were injected with a mixture of AAV1-Syn-cre-WPRE and AAV9-flex-tdTomato in LHA, and with AAV2/7-EF1a-DIO-hChR2(H134R)-eYFP in DRN. Alternatively, the mice were injected with a mixture of AAV1-Syn-cre-WPRE and AAV2/retro-CAG-flpO-P2A-TagBFP2 in LHA, and with a mixture of AAV2/9-EF1a-fDIO-hChR2(H134R)-eYFP and AAV9-CAG-flex-jrGECO in DRN. For all the electrophysiological experiments, the acute brain slices were prepared 3-weeks later for whole-cell patch clamp recordings.

For behavioral experiments with chronic silencing using tetanus toxin light chain (TeLC), the mice were bilaterally injected with a mixture of AAV1-hSyn-cre-WPRE an AAV2/9-CAG-Flex-tdTomato in the LHA, and AAV2/9-EF1a-DIO-eYFP (control) or AAV5-hSyn-Flex-TeLC-P2A-eYFP in DRN.

The coordinates for the injections were as follows: LH : −1.15 mm AP, ±1.0 mm ML, −5.2 mm DV; DRN : −4.65 mm AP, 0.0 mm ML, −3.35 and –3.4 mm DV.

### Electrophysiology

Three weeks after the injection of the virus carrying the channelrhodopin, the mice were anesthetized with isoflurane and were perfused transcardially with 10 mL of ice-cold NMDG-artificial cerebrospinal fluid (aCSF) solution containing (in mM): 1.25 NaH2PO4, 2.5 KCl, 10 MgCl2, 20 HEPES, 0.5 CaCl2, 24 NaHCO3, 8 D-glucose, 5 L-ascorbate, 3 Na-pyruvate, 2 thiourea, and 93 NMDG (osmolarity was adjusted to 300–310 mOsmol/L with sucrose). The pH was adjusted to 7.4 using 10 N HCl. Kynurenic acid (2 mM) was added to the perfusion solution on the day of the experiment. The brains were then quickly removed, and 250 μm acute brain slices encompassing the DRN were prepared using a Leica VT1200S vibratome. The slices were placed in a 32°C oxygenated perfusion solution for 10 min and were then incubated for 1 h at room temperature in HEPES-aCSF solution (in mM): 1.25 NaH2PO4, 2.5 KCl, 10 MgCl2, 20 HEPES, 0.5 CaCl2, 24 NaHCO3, 2.5 D-glucose, 5 L-ascorbate, 1 Na-pyruvate, 2 thiourea, 92 NaCl, and 20 sucrose (osmolarity was adjusted to 300–310 mOsmol/L with sucrose). The pH was adjusted to 7.4 using 10 N HCl. They were then transferred to a recording chamber on the stage of an upright microscope (Zeiss) where they were perfused with 3-4 mL/min of aCSF (in mM): 120 NaCl, 5 HEPES, 2.5 KCl, 1.2 NaH2P04, 2 MgCl2, 2 CaCl2, 2.5 glucose, 24 NaHCO3, and 7.5 sucrose). The perfusion chamber and the aCSF were kept at 32°C. All the solutions were oxygenated with 95% O2/5% CO2. A 60× water immersion objective and a video camera (Zeiss) were used to visualize neurons in the DRN. Borosilicate glass (3-5 MΩ resistance) recording pipettes were pulled using a P-1000 Flaming/Brown micropipette puller (Sutter Instruments). Recordings were performed using an Axopatch 200A amplifier (Molecular Devices). For the voltage-clamp recordings, the intracellular solution consisted of (in mM): 115 cesium methanesulfonate, 20 cesium chloride, 10 HEPES, 2.5 MgCl2, 4 Na2ATP, 0.4 Na3GTP, 10 Na-phosphocreatine, 0.6 EGTA, and 5 QX314, as well as 0.2% biocytin (pH 7.35). Signals were filtered at 5 kHz using a Digidata 1500A data acquisition interface (Molecular Devices, San Jose, CA) and acquired using pClamp 10.6 software (Molecular Devices). Pipette and cell capacitance were fully compensated. To examine monosynaptic transmission, the extracellular recording solution was supplemented with 1 μM TTX and 100 μM 4-AP. For the voltage-clamp experiments, postsynaptic currents were measured in DRN neurons clamped at −60 mV and 0 mV holding voltage following optogenetic stimulation of LH axon terminals with 5-ms blue light pulses delivered through the objective with a Colibri 7 LED light source (Zeiss). Excitatory and inhibitory transmissions were blocked with 3 mM NBQX and 10 mM gabazine, which are AMPA and GABA-a receptor antagonists, respectively. We calculated the excitation to inhibition ratio (E/I ratio) as a maximum amplitude recorded at –60 mV divided by the maximum amplitude recorded at 0 mV. E/I ratio values above 1 indicate larger contribution of excitatory component, and below 1 indicate larger contribution of inhibitory component. Once the recordings were completed, the slices were fixed in 4% formaldehyde for 24h and were then transferred to a 0.1M phosphate buffer solution for post hoc histological analysis (anti-tryptophan hydroxylase staining, described below). Spontaneous excitatory postsynaptic currents (sEPSCs) were recorded at −60 mV, and spontaneous inhibitory postsynaptic currents (sIPSCs) at 0 mV in gap-free mode for 5 min. sEPSC and IPSC frequency and amplitude were analyzed with Minhee Analysis Package^45^. The E/I ratio was calculated by dividing the frequency/amplitudes of EPSCs by the frequency/amplitudes of IPSCs from the same cell.

### Histology and immunostaining

The mice were deeply anesthetized using a mix of ketamine/xylazine (100 and 10 mg/kg, respectively, intraperitoneally) and were transcardially perfused with saline followed by a 0.1 M phosphate buffer solution (PB, pH7.4) containing 4% paraformaldehyde. The brains were postfixed overnight in the same solution, rinsed with PB, and stored in PB. Brain sections (100 μm for histology and 50 μm for immunostaining) were cut with a vibratome along the coronal plane.

Sections encompassing LHA were examined to confirm the accuracy of the injection sites. DRN sections were stained for TPH (tryptophan hydroxylase, a marker for 5-HT neurons) using a standard 2-day immunostaining protocol. Briefly, free-floating slices were first blocked in PB containing 5% normal donkey serum (NDS), 3% bovine serum albumine (BSA) and 0.4% Triton X-100 for 1.5 h. They were then incubated overnight with primary antibodies diluted in PB containing 3% BSA, 5% NDS and 0.4% Triton X-100 and then with a secondary antibody diluted in PB containing 3% BSA overnight. The primary antibodies were anti-TPH (Millipore, sheep polyclonal, 1:1000 dilution). The secondary antibody was donkey anti-sheep IgG AlexaFluor 647 (ThermoFisher Scientific, 1:1000 dilution). Sections from the electrophysiological recordings were immunostained against TPH using the protocol described above. Sections collected from the TeLC-injected mice were immunostained against TPH and cFos using the protocol described above. Immunostained sections were mounted using FluoromountTM Aqueous Mounting Medium (Millipore-Sigma)

### Imaging and image quantification

Confocal imaging was performed for all the visualizations in this study. Images were obtained from Zeiss LSM700 confocal microscope. Images were analyzed using ImageJ software. Cells from tracing experiments were manually counted using the plugin Colocalization Object Counter from Xmm^3 volume (or change to thickness). Images from cFOS quantification experiment were flatted using maximum intensity projection function, automatically segmented using plugin Trainable Weka Segmentation and automatically quantified using Colocalization Object Counter.

### Mouse Tissue Collection, Nuclei Isolation and Fixation

Mouse brains (N=6) were rapidly removed and dissected. To prevent RNA degradation, the brain tissue was immediately transferred to ice-cold PBS in centrifuge tubes and stored at − 80°C until processed for nuclei isolation.

Nuclei were first isolated through adaptations of previous published protocols^46^. Frozen tissue was briefly thawed on wet ice. The tissue sections were then transferred to chilled lysis buffer (250 mM sucrose, 25 mM KCl, 5 mM MgCl2, 10 mM Tris-HCl, 1 µM DTT, 0.4 U/µl Enzymatic RNase-In, 0.2 U/µl Superase-In, DNAse Inhibitor (1u/million cells) 0.1% Triton X-100), dissociated until homogeneous with a pestle, and incubated on ice for 10 minutes. The nuclei suspension was passed through a 70 µm cell strainer and centrifuged at 500xg for 5 minutes at 4°C to pellet the nuclei. The nuclei pellets were resuspended in 750 µL nuclei buffer containing 0.75% BSA (Parse Biosciences) and then fixed using the Parse Biosciences Nuclei Fixation kit (Evercode Nuclei Fixation v2). Briefly, samples were passed through a 40 µm filter, incubated in the nuclei fixation solution for 10 minutes on ice, followed by incubation in the nuclei permeabilization solution. Nuclei neutralization buffer was added to each sample, and centrifuged at 500xg for 10 minutes at 4°C. The pellets were resuspended in nuclei buffer with DMSO (1:20), frozen, and stored at −80°C until library preparation.

#### Barcoding and Library Preparation

Fixed nuclei were removed from −80°C and thawed in a water bath set to 37°C, placed on ice, and counted. Barcoded single-cell libraries were prepared from fixed single-cell suspensions using Evercode Whole Transcriptome v.2 (Parse Biosciences) following the manufacturer’s instructions. Briefly, fixed nuclei underwent three rounds of barcoding in three reaction plates to generate single-nucleus RNASeq libraries. In the first round, nucleus-specific barcodes were added through an *in situ* reverse transcription. At the end of the reaction, nuclei from all wells of the Round 1 plate were pooled and combined with a ligation mix. The nuclei suspension was then added to each well of the Round 2 barcoding plate and incubated. After stopping, the Round 2 ligation nuclei were pooled and filtered through a 40 µm strainer. The filtered nuclei underwent a third round of barcoding by ligation in the Round 3 plate. After adding the Round 3 stop mix, the nuclei were pooled again, filtered through a 40 µm strainer, washed, and counted. The pooled nuclei were distributed into multiple sub-libraries based on the manufacturer’s instructions. cDNA isolated from each sub-library was then used to generate an Illumina-compatible sequencing library.

#### Single-Nuclei RNA Sequencing

Sub-libraries were pooled and sequenced on an Illumina NovaSeq 6000 platform (Plateforme de Séquençage de Nouvelle Génération (NGS), CRCHU de Québec (CHUL)) with a sequencing configuration of paired-end 100 bp, generating 50,000 reads per nucleus.

### Bioinformatic analyses

#### Quality check, alignment and feature quantification

Firstly, using the M33 release of the GRCm39 mouse genome, we built a reference genome with the “mkref” mode of the Parse Biosciences Pipeline (version1.1.2). We added into the genome the transgene-specific sequences of the viral strategies (**Table S3**) used in our experiments. The pipeline which performs quality check (FastQC), demultiplexing, aligning (STAR) and feature quantification, was ran for each sub-library separately and then, in “combine” mode with the whole dataset to produce a unique count matrix.

#### Standardization, clustering and visualization

We kept barcodes (from now on considered as cells) with a transcript count between 200 and 400000, that expressed at least 100 unique genes and for which the mitochondrial gene expression percentage was less than 5%. This resulted in a total of 1832 high-quality nuclei that passed these thresholds. The 1832-cell dataset was normalized using SCTtransform, which applies regularized negative binomial regression to regress out technical covariates, including mitochondrial gene percentage, and to stabilize variance across cells^47^.

Following a standard Seurat pipeline, we initially identified the nearest 20 neighbors for each Pearson residual and identified clusters using the Leiden algorithm with varying resolutions (0.1,0.25,0.5,0.75 and 1). The cluster identity was assessed according to differential expression analysis (Wilcoxon rank sum test) and automatically annotated with the MapMyCells software (https://portal.brain-map.org/atlases-and-data/bkp/mapmycells). Based on established scRNA-seq annotation strategies^48^, and informed by prior DRN molecular classification^32^, cluster identity was manually curated using cluster size, hierarchal class and subclass annotations, canonical marker gene expression, and the proportion of cells expressing defining markers within each cluster.

#### Neuronal assessment and transfection labeling

Considering our sample size and a high risk of error type 1 by the number of comparisons, we reperformed the clustering steps keeping only the cells annotated as neurons. Within the latter, we assessed the expression of some non-neuronal marker genes to estimate a count threshold between signal and noise. Based on this, we labeled any cell that expressed more than 5 raw counts of EGFP or mCherry as transgene expressive. Finally, we performed a pseudo-bulk differential expression analysis with edgeR^49,50^ with the raw counts, filtering by expression (greater than 5 to be consistent) and comparing non-transgene expressing cells to those expressing one or both transgenes. The gene ontology terms enrichment in our set of DEGs was also evaluated using the R package g:Profiler2.

#### Serotoninergic assessment

Given the complex serotoninergic diversity of the DRN^1,32,33^ and the fact that distinct groups of serotonergic neurons were observable in the visualizations and the clustering, we decided to perform re-clustering steps keeping only the serotonergic neurons and describing the molecularly distinct groups.

The analyses were carried out in the R environment and python (MapMyCells). No sample size calculation was determined; multiple test corrections were done using Benjamini-Hochberg and the critical value α was set at 0.05 for hypothesis testing unless stated otherwise.

### Behavioral experiments

#### Open Field Test (OFT)

The experimental mice were placed in an open field arena (50 cm x 50 cm) for 10 minutes. The arena was novel for animals. The experiments were conducted in the dim light (around 80 lux). Center point of the mouse was tracked using a web camera LoGiTech and ANY-maze video-tracking software. The arena was virtually divided into inner (center) and outer zones as described in Seibenhener et al.^51^. Mobility score was quantified using AnyMaze software based on frame-to-frame changes in pixel intensity. Rotations of the animal’s body were quantified using AnyMaze software as the number of full 360° rotations. The animal’s center point was defined as a virtual origin that remained constant across frames. A vector was constructed from the center point to the head position, and the angle between successive vectors was calculated. As long as the direction of rotation remained consistent (i.e., the angle retained the same sign), angular changes were accumulated. A full rotation was defined when the cumulative angle reached 360°.

#### Novelty Suppressed feeding test (NSFT)

The NSFT was conducted as previously described^52^. 24h prior to the test experimental mice were food deprived. The day of the experiment, the mice were placed in an open field arena (50 cm x 50 cm) which had a different wall color and scent from one used in OFT. This arena was novel for animals. Animals’ behaviour was recorded and tracked as described above. In the center of the arena we placed one food palette. The experiments were conducted in the red light. Time from the start of the experiment to the moment of chewing on the food palette was estimated and further referred to as ‘latency to eat’.

#### Nestlet shredding test

Test was conducted as described by Angoa-Perez et al.^53^. Briefly, experimental mice were separated and single-caged for 1 h in the new cages, containing new intact nest material. Prior to testing, the mice were acclimated to the testing room for at least 30 minutes. The experiments were conducted in the dim light (around 80 lux).

#### Tail suspension test (TST)

TST was conducted as previously detailed^27^. Briefly, mice were suspended by their tail for 10 min in the custom arena, and were tracked the same way as in OFT. As this test is aversive and stressful for the animals, it was performed as a last one before single-caging the experimental mice. TST was reperformed before sacrificing the animals for post hoc histological analysis of cFos expression.

#### Marble burying test

The experimental setup involved an open-topped plastic cage (dimensions: [insert dimensions if known],) filled with a Sani-Chip bedding material to a depth of approximately 5 cm. The bedding was evenly spread to provide a uniform substrate for burying. The test cage contained 20 glass marbles, each with a diameter of approximately 1.5 cm, placed in an evenly spaced 4 x 5 grid pattern on the surface of the bedding. The experiments were conducted in the dim light (approx. 80 lux). Prior to testing, the mice were acclimated to the testing room for at least 30 minutes. Then, each mouse was individually placed into the test cage and allowed to explore freely for a period of 1h. During this time, the animal could interact with the marbles and display natural burying behaviors. After the testing period, the number of marbles buried, defined as at least two-thirds covered by bedding, was recorded.

#### Splash test

The behavior test was conducted as detailed before^52^. A 10% sucrose solution was prepared in MilliQ water, which acts as a mildly sticky substance that prompts grooming behavior in mice. The experimental setup comprised a clean, open-topped cage. The cage was free of bedding to provide a clear view of grooming behavior during testing. The experiments were conducted in the dim light (approx. 80 lux). Prior to testing, the mice were acclimated to the testing room for at least 30 minutes. Two sprays (typically 0.1-0.2 mL) of the 10% sucrose solution were applied directly to the dorsal coat of each mouse using a spray bottle. Immediately after applying the sucrose solution, each mouse was placed into the cage for a period of 5 minutes. The cumulative time spent grooming within the 5-minute observation period was counted manually from the recorded videos.

#### Resident-intruder test

The test was conducted as previously described by Koolhaas et al.^54^. Experimental animals were removed from group housing and singly housed for one week prior to testing. The experiment was performed in the home cage under dim light conditions (approximately 80 lux) and lasted 10 minutes.

#### Sucrose preference test

In their home cages, mice were habituated to a two-bottle choice setup for 24 hours before testing, both initially containing water. Next day, one bottle was replaced with a bottle containing the 2% sucrose solution. The bottles were identical and positioned side-by-side to prevent any spatial bias. The position of the bottles was switched on the second day of testing to avoid side preference. The liquid consumption was measured daily, and eventually the sucrose preference score was calculated as a percentage of sucrose solution consumed out of overall liquid intake.

#### Pellet consumption

Mice were singly housed and had free access to food pellets. Pellet consumption was measured by tracking changes in pellet weight over three days and calculating the average consumption across this period.

#### Social preference test

The test was conducted as described elsewhere^55^. Briefly, the experiment took place in an open-field arena equipped with two small wire cages positioned on opposite sides. Each experimental animal underwent four stages: habituation to the apparatus, habituation to the presence of objects in the cages, the social preference test, and the social novelty test. All experiments were performed under dim lighting conditions (∼80 lux).

### Data and statistical analysis

Data analysis, statistical analyses, and data visualization were performed using Python and R. Electrophysiological data were analyzed using the Neo IO Python package and custom scripts. Linear and linear mixed-effects models were applied to raw or log-transformed data that met normality assumptions. Count data were analyzed using negative binomial or binomial generalized linear models. When appropriate, post hoc pairwise comparisons were performed on estimated marginal means using Tukey adjustment for multiple comparisons. Most models used for Figures 5 and 6 were adjusted for sex and batch effects. Statistical significance was set at p < 0.05. The specific statistical tests used are indicated in the figure legends, and detailed information for each analysis is provided in **Table S1**.

#### Sample size

The sample size for each group for behavioral, anatomical, and electrophysiological experiments was determined from previously published work and from pilot experiments performed in our laboratory.

