## Supplemental Figures for "Lateral hypothalamic input engages a disinhibitory microcircuit in the dorsal raphe to promote behavior activation"

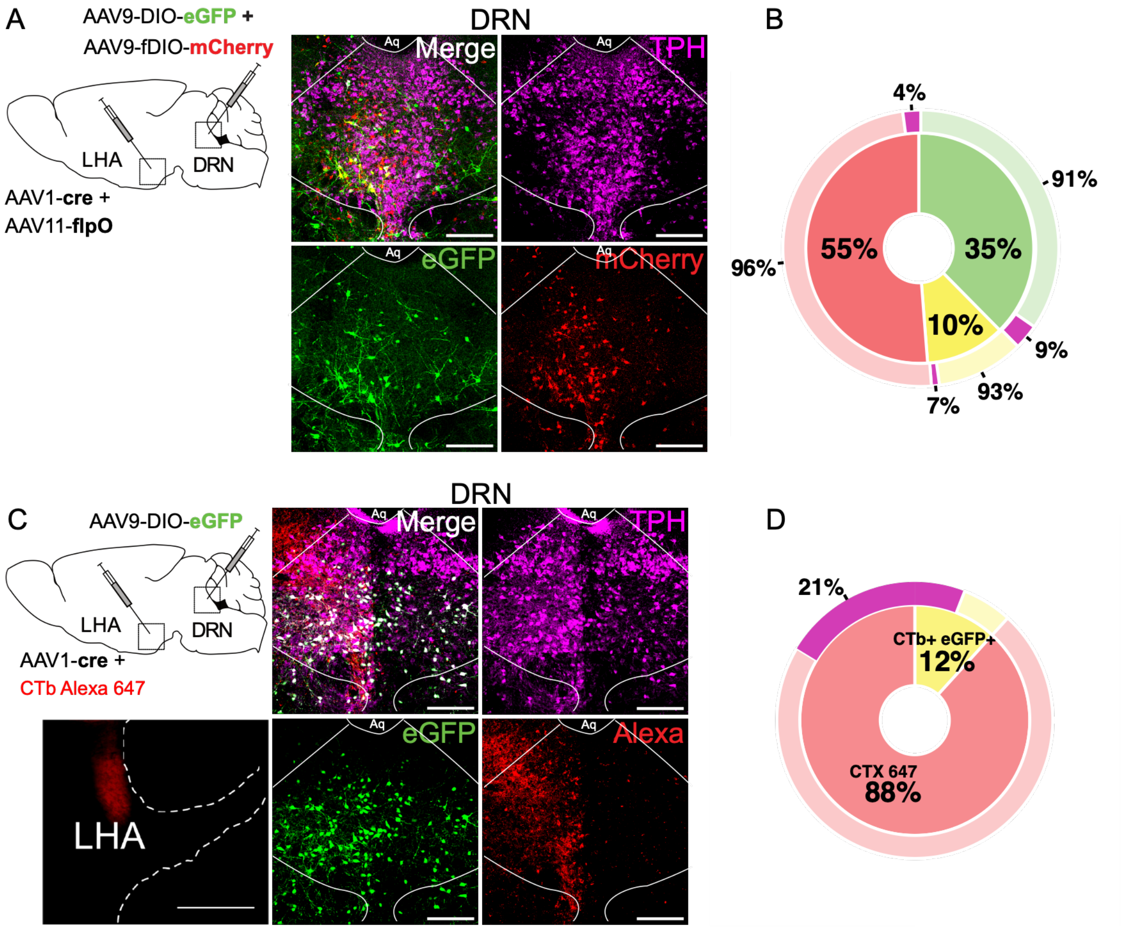


**Figure S1. Alternative retrograde tracers confirm limited overlap between LHA-innervated and LHA-projecting DRN neurons**

A. Schematic of the dual-labeling strategy combining anterograde transsynaptic labeling of DRN neurons receiving LHA inputs (eGFP+) with retrograde labeling of DRN→LHA projection neurons using AAV11-FlpO and Flp-dependent mCherry expression. Representative DRN images show eGFP (green), mCherry (red), and TPH immunoreactivity (magenta).

B. Quantification of eGFP+, mCherry+, and double-labeled (eGFP+ mCherry+) DRN neurons (n = 3 mice, 2565 cells). Outer ring indicates the fraction of TPH+ neurons within each category (magenta).

C. Schematic of the strategy combining eGFP labeling of DRN neurons receiving LHA inputs (eGFP+) with retrograde labeling of DRN→LHA neurons using CTb-Alexa$^{647}$. Representative DRN images show eGFP (green), CTb-Alexa$^{647}$ (red), and TPH immunoreactivity (magenta), including the LHA injection site.

D. Quantification of CTb-Alexa$^{647}$ and double-labeled (eGFP+ CTb-Alexa$^{647}$) DRN neurons (n = 4 mice, 748 cells) Outer ring indicates the fraction of TPH+ neurons within each category (magenta).

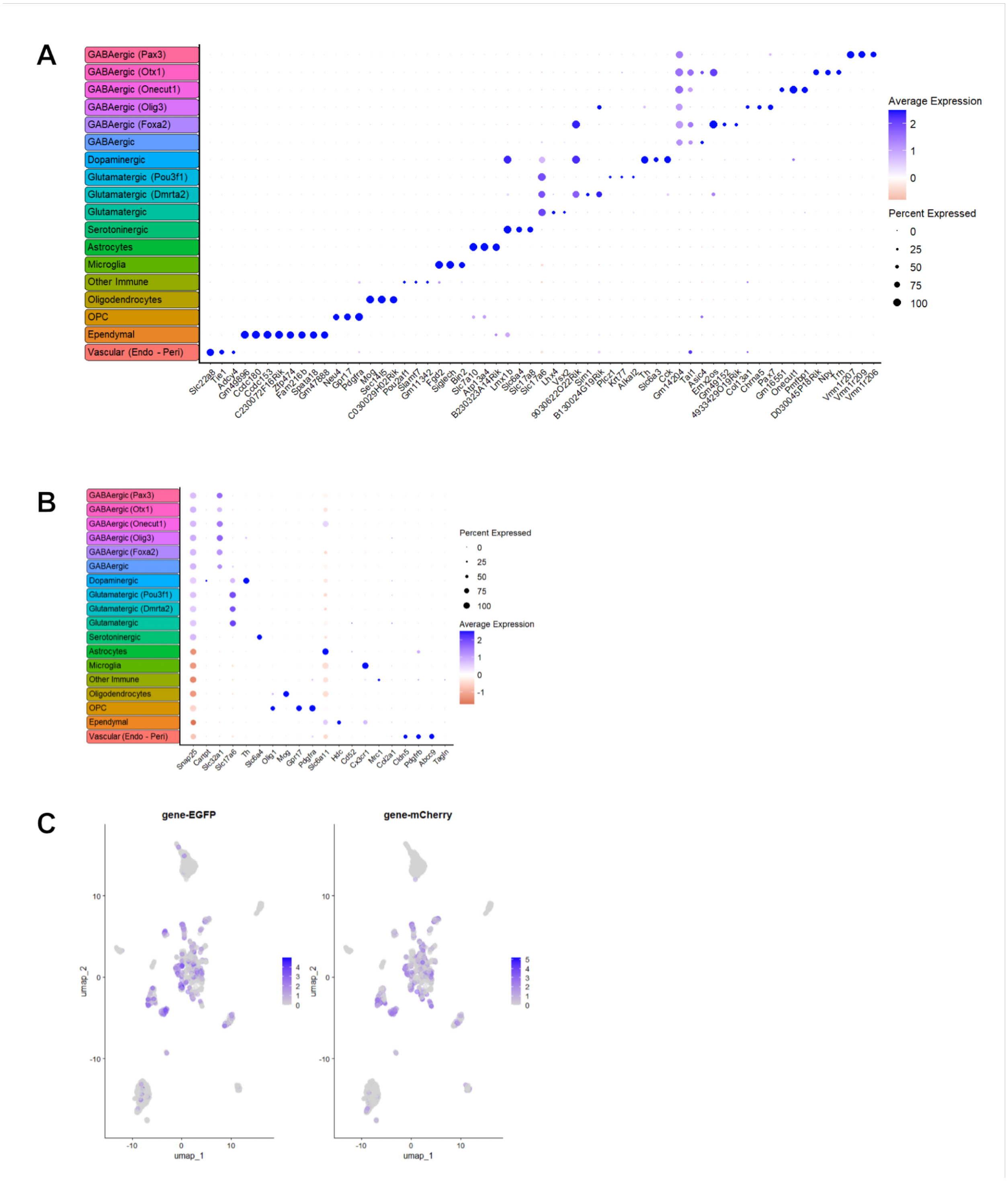


**Figure S2. Validation of DRN cell-type annotation and transgene detection in the snRNA-seq dataset**

A. Dot plot showing the top differentially expressed markers genes for each annotated DRN cell type identified in our dataset. Differential expression was assessed using

Wilcoxon rank-sum tests comparing each cluster to all others. Dot size represents the percentage of cells expressing the gene within cluster; color indicates scales average expression (SCT-normalized counts).

B. Dot plot of previously reported DRN marker genes (Huang et al. 2019) projected onto our dataset. Differential expression was assessed using Wilcoxon rank-sum tests for each cluster versus the remainder of the dataset. Dot size represents the percentage of cells expressing the gene; color represent the scaled average expression.

C. UMAP projection showing normalized expression of the transgene transcripts (gene-eGFP and gene-mCherry) across neurons in the dataset.

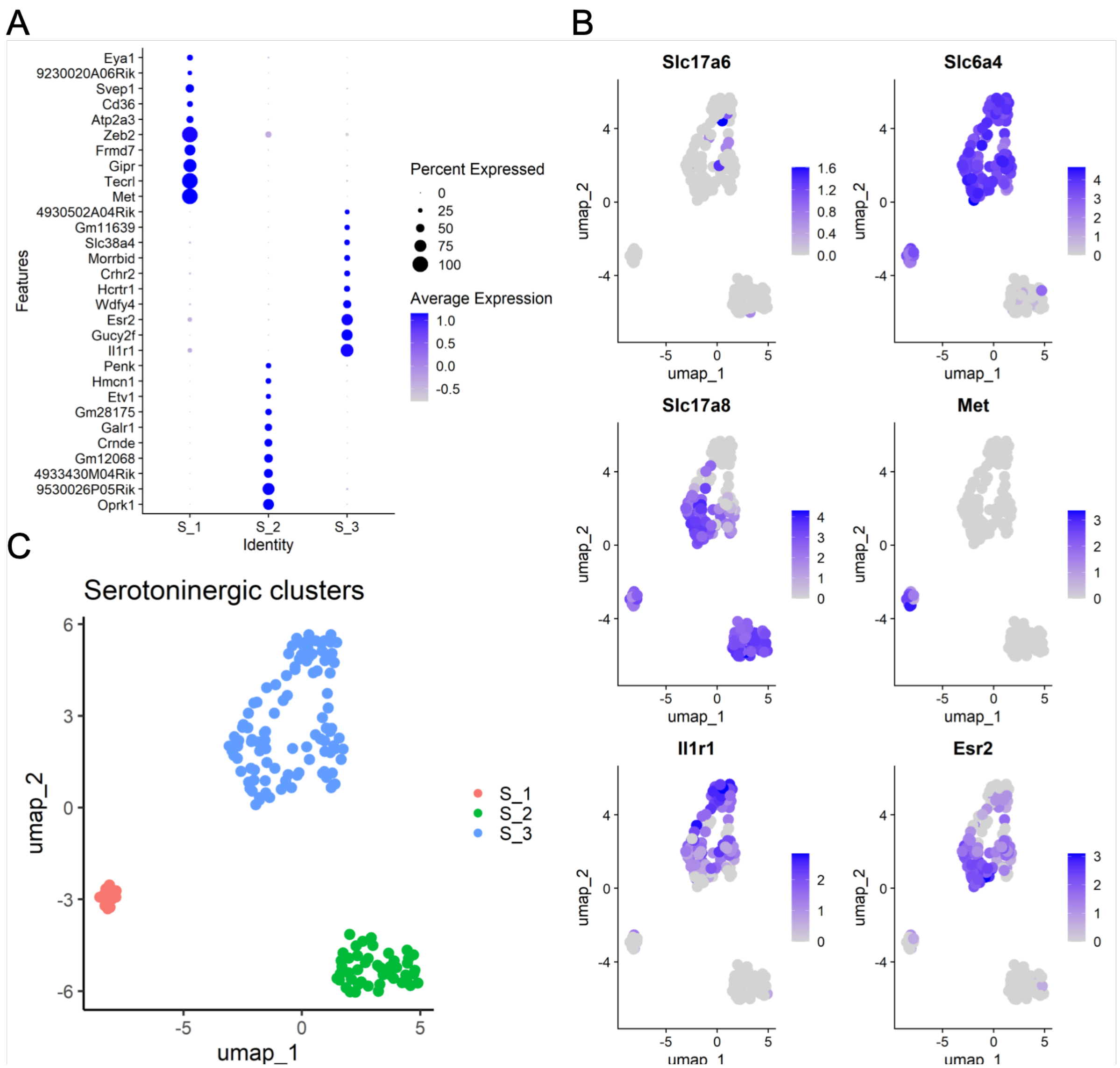


### Figure S3. Transcriptional heterogeneity of serotonin DRN neurons

A. Dot plot showing the top differentially expressed markers genes across the three serotonergic neuron clusters identified in the dataset (S_1, S_2, S_3). Differential expression was assessed using Wilcoxon rank-sum test comparing each cluster to the remaining serotonergic cells. Dot size represents the percentage of cells expressing the gene; color indicates scales average expression (SCT-normalized counts).

B. UMAP projections showing the normalized expression expression of representative differentially expressed genes distinguishing serotonergic clusters (e.g., *Slc17a6*, *Slc6a4*, *Il1r1*, *Esr2*, *Met*). Color scale reflects normalized expression levels.

C. UMAP projection of serotonergic neurons showing the spatial distribution and relative size of the three transcriptionally defined clusters (S_1, S_2, S_3)..

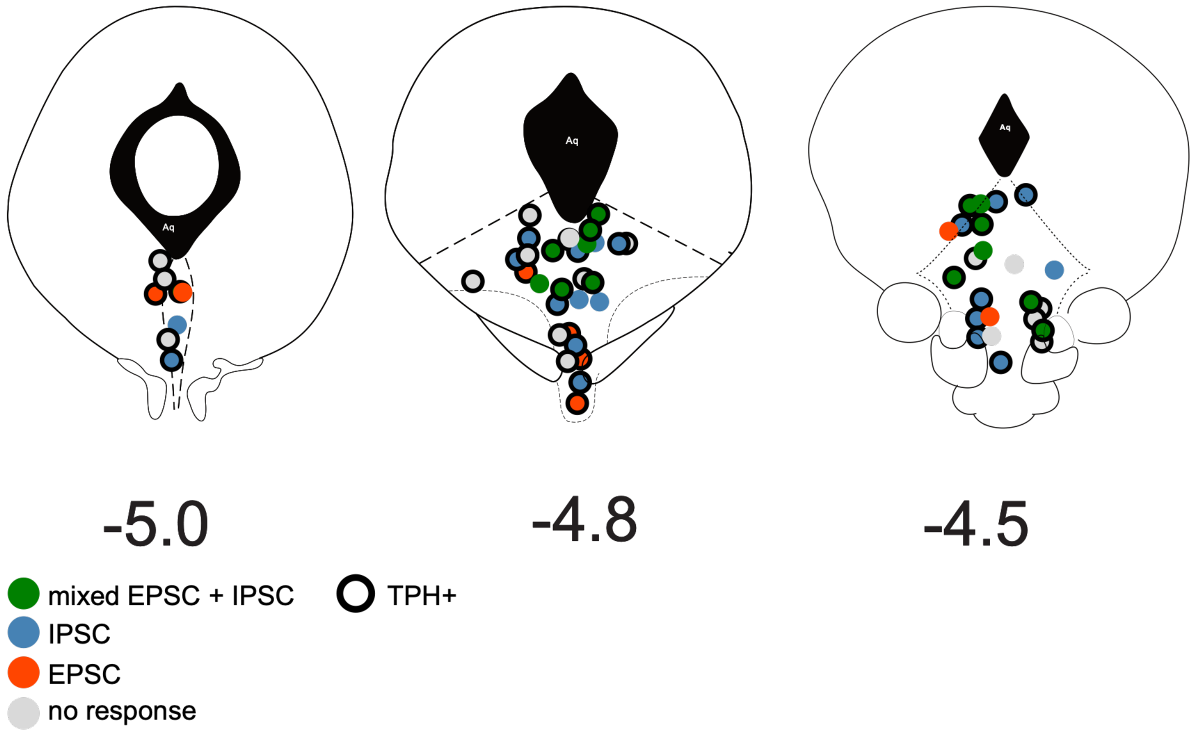


**Figure S4. Recording locations and synaptic response of DRN neurons analyzed in Figure 3**

Schematic coronal representations of the dorsal raphe nucleus (DRN) at three rostrocaudal levels (−5.0, −4.8, and −4.5 mm from bregma) showing the spatial distribution of neurons recorded in Figure 3. Each circle represents one recorded neuron and is color-coded by synaptic response to optogenetic stimulation of LHA terminals: IPSC only (blue), EPSC only (orange), mixed EPSC+IPSC (green), or no detectable response (gray). Cells outlined in black were identified as serotonergic (TPH+). Aq, cerebral aqueduct.

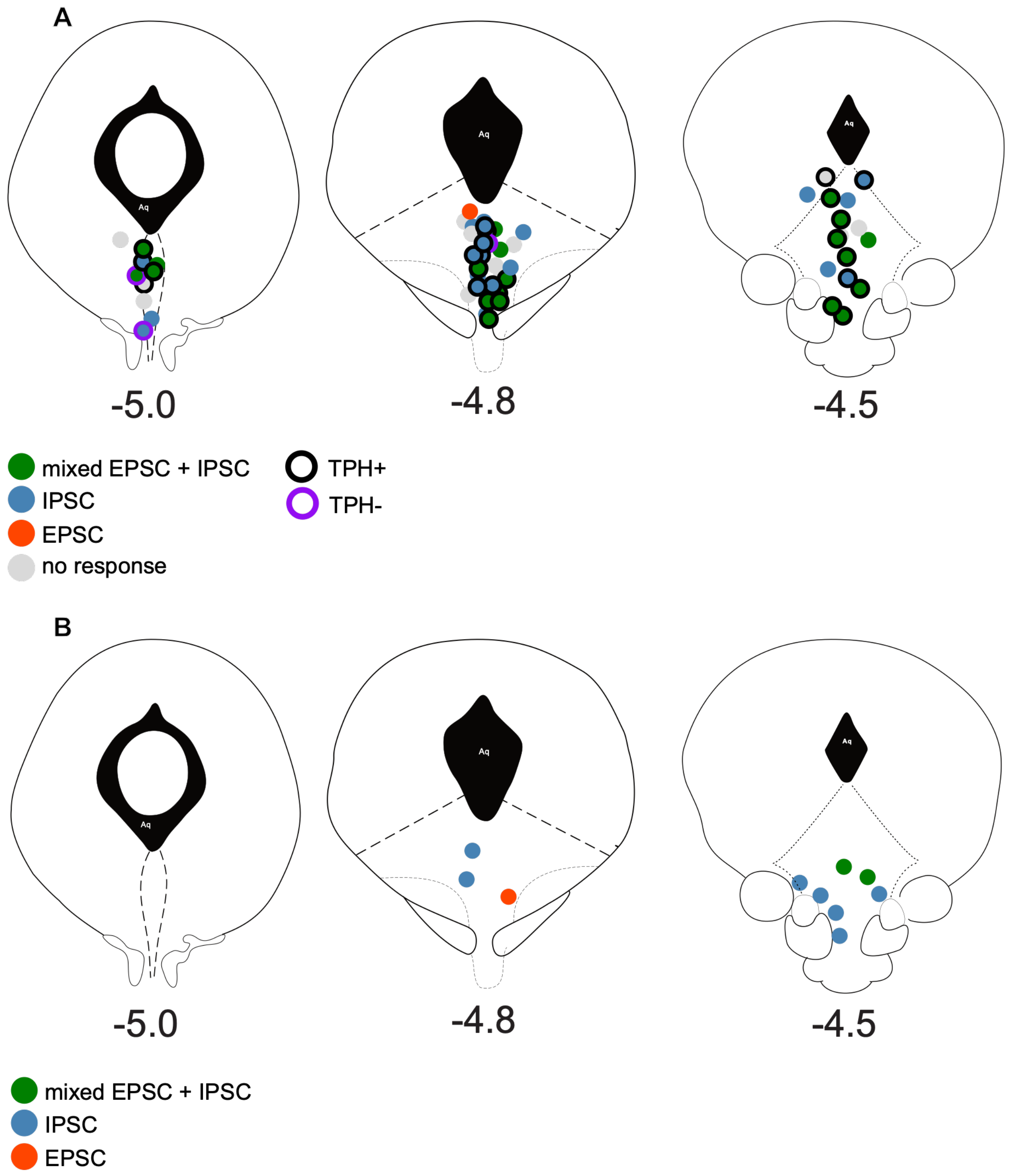


**Figure S5. Recording locations and synaptic responses for intra-DRN connectivity experiments (Figure 4)**

Schematic coronal representations of the DRN at three rostrocaudal levels (−5.0, −4.8, and −4.5 mm from bregma) showing the spatial distribution of neurons recorded in Figure 4.

A. Recordings from DRN neurons during optogenetic stimulation of LHA-innervated DRN neurons. Each circle represents one recorded neuron and is color-coded by synaptic response class: IPSC only (blue), EPSC only (orange), mixed EPSC+IPSC (green), or no detectable response (gray). Serotonergic identity is indicated by black outline (TPH+) or magenta outline (TPH−).

B. Recordings from DRN neurons during optogenetic stimulation of DRN→LHA projection neurons. Circles are color-coded by synaptic response class as in (A). Aq, cerebral aqueduct.

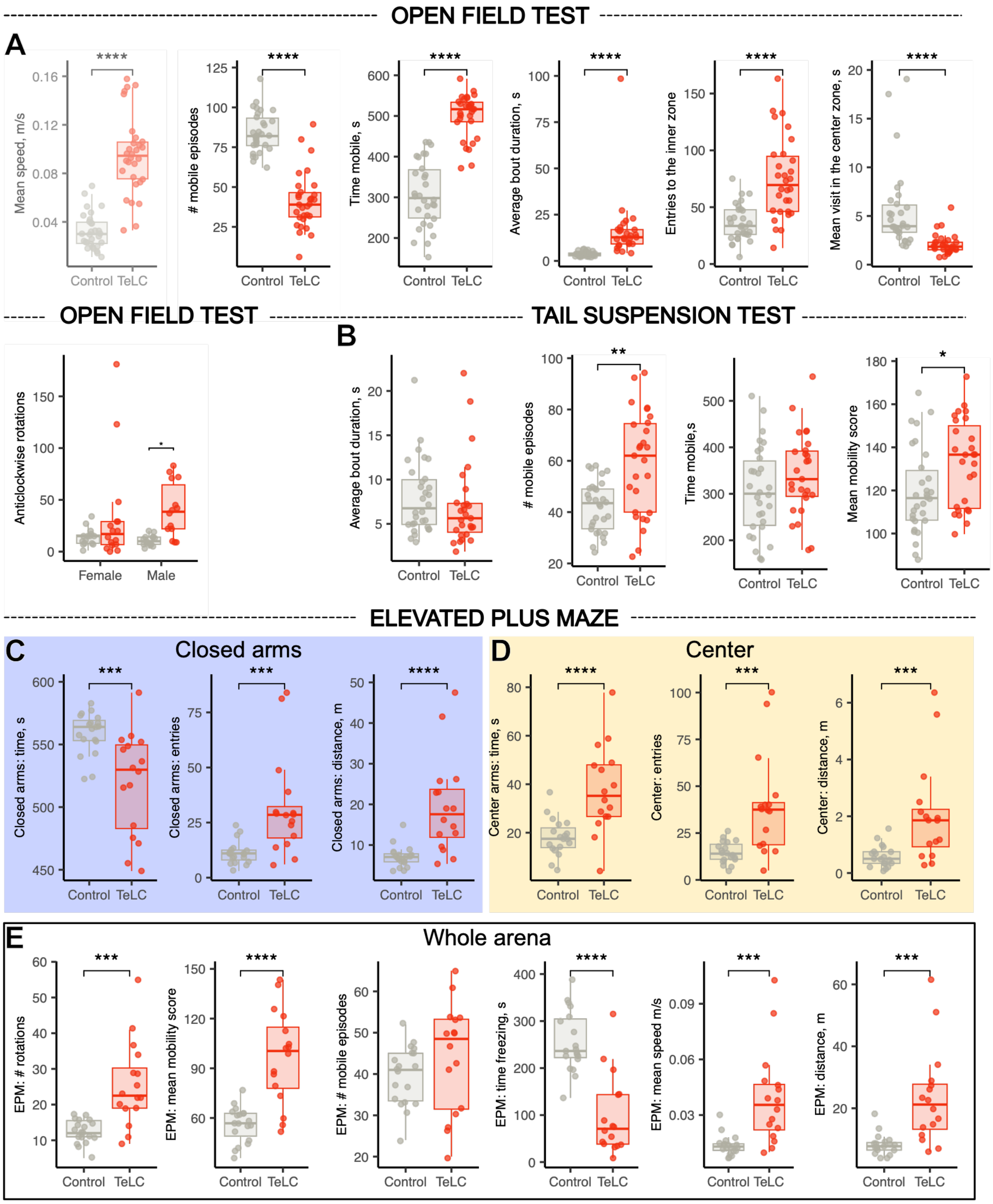


**Figure S6. Expanded locomotor and mobility analyses following chronic silencing of LHA-tergeted DRN neurons**

A. Open field test (OFT): detailed locomotor parameters including mean speed, number of mobile episodes, total time mobile, average bout duration, number of entries to the center zone, and mean visit duration in the center, OFT rotation analysis separated by sex (anticlockwise rotations).

B. Tail suspension test (TST): average bout duration, number of mobile episodes, total time mobile, and mean mobility score.

C-E. Elevated plus maze (EPM) detailed locomotor and zone-specific metrics.

C. Closed-arm parameters: time spent, number of entries, and distance traveled.

D. Center zone parameters: time spent, number of entries, and distance traveled.

E. Whole-arena measures: total rotations, mean mobility score, number of mobile episodes, time freezing, mean speed, and total distance traveled.

Boxplots show median and interquartile range (IQR), with whiskers extending to 1.5× IQR; individual points represent mice. Significance levels are indicated in panels. Statistical models correspond to those described for Figure 5. *$p < 0.05$, **$p < 0.01$, ***$p < 0.001$ and ****$p < 0.0001$.

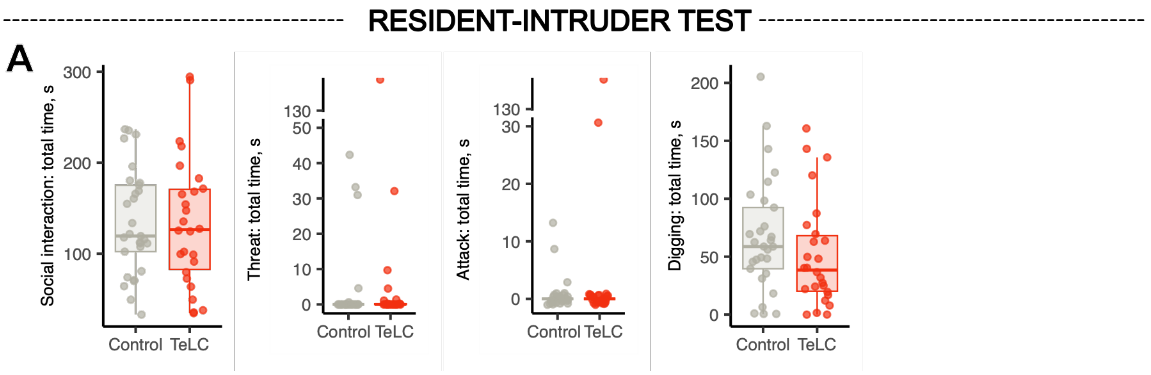


**Figure S7. Chronic silencing of LHA-targeted DRN neurons does not alter social interaction or aggression in the resident-intruder test**

A. Quantification of behavioral parameters in the resident–intruder assay, including total time spent in social interaction, threat behavior, attack behavior, and digging. No significant differences were detected between control and TeLC-expressing mice across measured parameters.

Boxplots show median and interquartile range (IQR), with whiskers extending to 1.5× IQR; individual points represent mice. Statistical analyses correspond to those described for Figure 5.

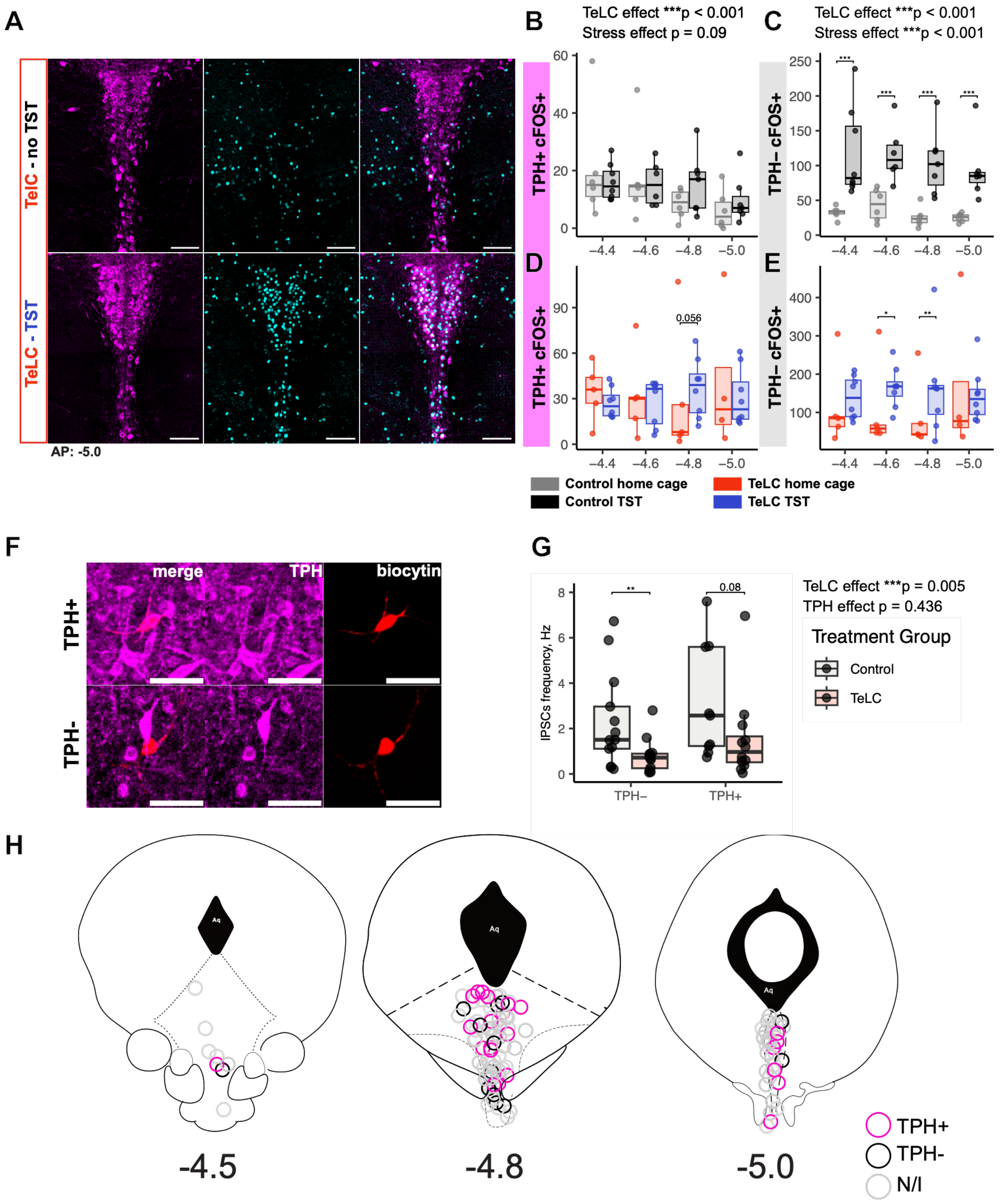


**Figure S8. Stress- and cell type-specific effects of TeLC-mediated silencing in the DRN**

**A.** Representative confocal images of DRN coronal sections at −5 mm from bregma (scale bar: 200 μm) showing cFos (cyan) and TPH (magenta) immunostaining in control (eYFP) and TeLC-expressing mice under basla (home cage) and acute stress (TST) conditions.

**B–C.** Effect of acute stress (TST; black) on the number of $cFos^{+}$ DRN neurons across the anterior–posterior axis in control YFP mice, shown separately for $TPH^{+}$ and $TPH^{-}$ populations. The x-axis represents distance from bregma (mm).

**D–E.** Effect of acute stress (TST; blue) on the number of $cFos^{+}$ DRN neurons across the anterior–posterior axis in TeLC-expressing mice, shown separately for $TPH^{+}$ and $TPH^{-}$ populations. The x-axis represents distance from bregma (mm).

**F.** Representative confocal image of biocytin-filled DRN neurons following whole-cell recordings, with post hoc TPH immunostaining. Examples are shown for $TPH^{+}$ and $TPH^{-}$ neurons.

**G.** Effect of TeLC expression on spontaneous IPSC (sIPSC) frequency in $TPH^{+}$ and $TPH^{-}$ DRN neurons. TeLC significantly reduced sIPSC frequency ($p = 0.005$), with no significant effect of TPH identity ($p = 0.436$) and no TeLC x TPH interaction ($p = 0.584$).

**H.** Anatomical distribution of recorded neurons across rostro-caudal DRN (−4.5, −4.8, −5.0 mm from bregma). Circles indicate TPH+ (magenta), TPH- (black), and neurons not identified following post hoc staining (N/I, gray).

Boxplots show median and interquartile range (IQR), with whiskers extending to 1.5× IQR; individual points represent cells or mice as indicated. Statistical details are provided in Supplementary Tables.
